# Furin controls β cell function via mTORC1 signaling

**DOI:** 10.1101/2020.04.09.027839

**Authors:** Bas Brouwers, Ilaria Coppola, Katlijn Vints, Bastian Dislich, Nathalie Jouvet, Leentje Van Lommel, Natalia V. Gounko, Lieven Thorrez, Frans Schuit, Stefan F. Lichtenthaler, Jennifer L. Estall, Jeroen Declercq, Bruno Ramos-Molina, John W.M. Creemers

## Abstract

Furin is a proprotein convertase (PC) responsible for proteolytic activation of a wide array of precursor proteins within the secretory pathway. It maps to the PRC1 locus, a type 2 diabetes susceptibility locus, yet its specific role in pancreatic β cells is largely unknown. The aim of this study was to determine the role of furin in glucose homeostasis. We show that furin is highly expressed in human islets, while PCs that potentially could provide redundancy are expressed at considerably lower levels. β cell-specific furin knockout (βfurKO) mice are glucose intolerant, due to smaller islets with lower insulin content and abnormal dense core secretory granule morphology. RNA expression analysis and differential proteomics on βfurKO islets revealed activation of Activating Transcription Factor 4 (ATF4), which was mediated by mammalian target of rapamycin C1 (mTORC1). βfurKO cells show impaired cleavage of the essential V-ATPase subunit Ac45, and by blocking this pump in β cells the mTORC1 pathway is activated. Furthermore, βfurKO cells show lack of insulin receptor cleavage and impaired response to insulin. Taken together, these results suggest a model of mTORC1-ATF4 hyperactivation in β cells lacking furin, which causes β cell dysfunction.

## Introduction

Many protein hormones are synthesized as large inactive precursors that need a controlled series of proteolytic events to be eventually secreted into the bloodstream as active hormones. Any alterations of this regulated process can result in pathological conditions. Typically, when this alteration includes pro-hormones involved in regulation of glucose and lipid homeostasis, it can result in metabolic disorders (1–3).

Proprotein convertases (PCs) are subtilisin-like serine proteases that cleave a variety of precursor proteins in the secretory pathway such as hormones, neuropeptides, extracellular matrix proteins, membrane receptors and their ligands (4). Several studies have established an important role for the proteolytic activity of certain PCs in metabolic disorders. For instance, PC1/3 and PC2 are both highly expressed in the secretory granules of neuronal and endocrine cells and process a number of hormones and neuropeptides involved in the regulation of glucose and energy homeostasis such as insulin, glucagon and proopiomelanocortin (POMC) (1, 2). In humans, rare homozygous and compound heterozygous mutations that cause complete loss of PC1/3 enzymatic activity lead to hyperphagic obesity and impaired glucose homeostasis, among other endocrinopathies (5). In addition, common heterozygous PC1/3 mutations that cause partial loss-of-function contribute to variation in human body mass index (BMI) and plasma proinsulin and are associated with impaired regulation of plasma glucose levels (6). On the other hand, genome-wide association studies have found an association between several common variants in the gene encoding human PC2 with diabetes risk and related traits in human populations (7–10).

In contrast, the role of the PC family members active in the constitutive secretory pathway, such as furin, PC5/6, PACE4 or PC7 in obesity and type 2 diabetes (T2D) has been much less explored. In the secretory pathway furin is active in the *trans*-Golgi network (TGN), endosomes and at the cell membrane. In neuroendocrine cells furin has also been found in immature secretory granules, where it has been proposed to participate in the processing of certain neuropeptides involved in energy metabolism, such as pro-brain-derived neurotrophic factor (BDNF), pro-growth hormone releasing hormone (GHRH) and pro-kisspeptin (KISS1) (11–13). Furin also cleaves certain semaphorins that control energy balance by acting as guidance molecules for melanocortin circuits (14). The *FURIN* gene is located near to the PRC1 T2D association interval and was therefore suggested to be a promising biological T2D candidate (15). In addition, polymorphisms in *FURIN* have been associated with the prevalence of metabolic syndrome (16). In pancreatic β cells, furin has been shown to be crucial for acidification of secretory granules via the cleavage of Ac45, an essential subunit of the V-ATPase proton pump (17). The V-ATPase complex acidifies intracellular organelles by pumping protons across membranes, and acidification is a critical step in β cell granule maturation and proinsulin-to-insulin conversion by PC1/3 and PC2 (18). However, the significance of protein processing by furin in β cell function is incompletely understood.

In this study we have examined the expression levels of *FURIN* and other PCs in islets of healthy individuals and T2D patients. Furthermore, we have established the role of furin in β cells by creating a conditional knockout mouse model and knockout β cell lines. Metabolic studies were performed to characterize the function of furin in pancreatic β cells *in vivo*. Genomics and proteomics studies allowed to determine pathways affected by the absence of furin and to identify physiological furin substrates that influence β cell function.

## Results

### PC expression in human islets of healthy and T2D donors

In order to determine the potential importance of furin in human islets, we first quantified mRNA expression levels of *FURIN* and other PCs in islets from healthy and T2D donors (**Figure 1)**. *PC1/3* and *PC2*, which are responsible for processing of proinsulin, proglucagon and prosomatostatin are, as expected, expressed highest. *FURIN* showed highest expression of the PCs mainly active in the constitutive secretory and endosomal pathway (*PC5/6*, *PACE4*, and *PC7*), which is consistent with previously published RNAseq datasets from human islets and islet subsets (19, 20) (**Figure S1A-B**). Expression levels of these PCs that can potentially provide redundancy for furin activity are expressed at much lower levels, which could point to a non-redundant function of furin in human β cells. Except for slight increases in *PC5/6* and *PACE4* expression, we did not observe significant changes in PC expression in islets from T2D donors. *INSULIN* expression was significantly reduced in T2D patients in this cohort (**Figure 1**).

**Figure 1.**
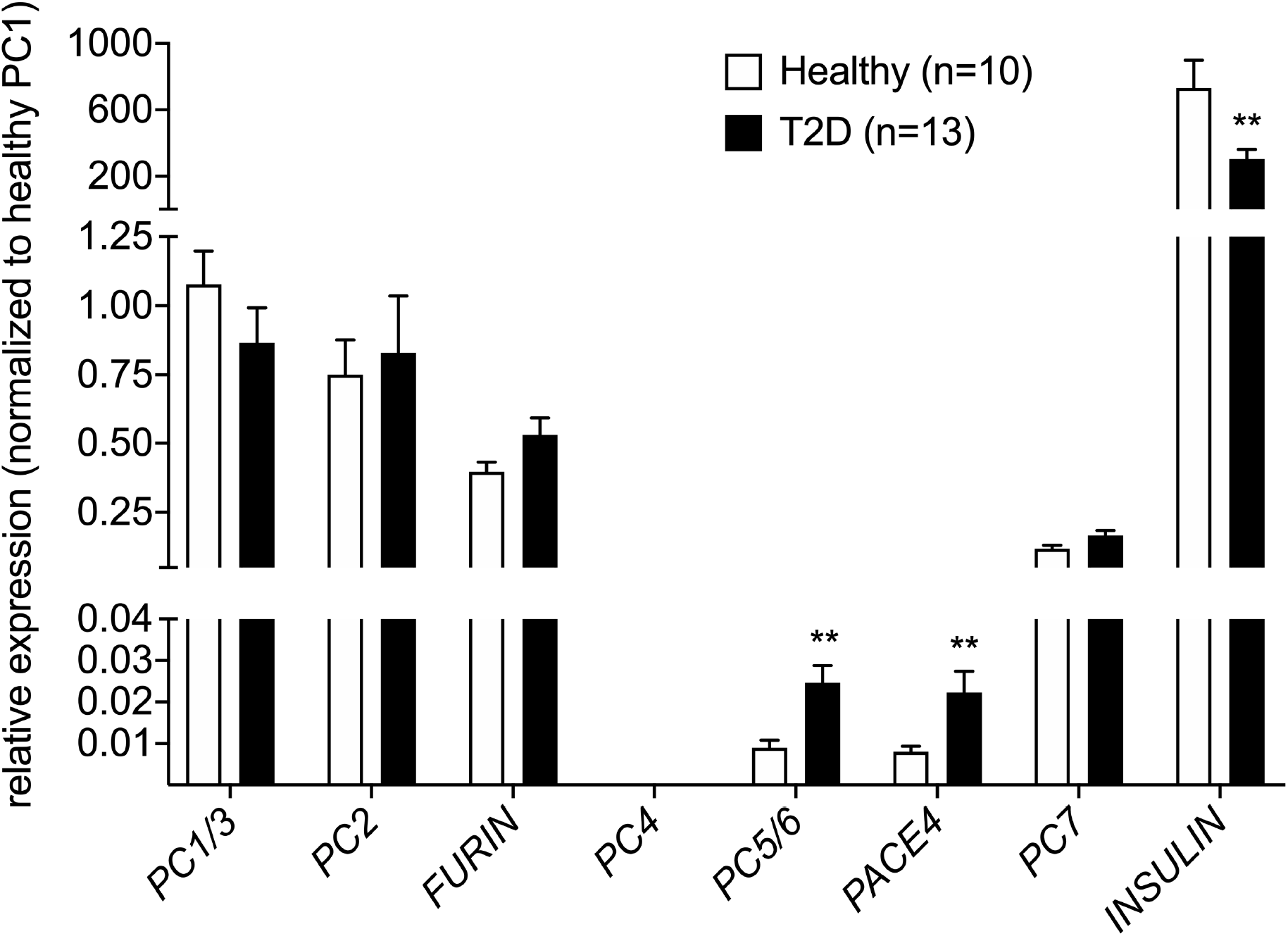
Furin is highly expressed in human islets. qRT-PCR analysis of 7 closely related PCs in islets from healthy (n=10) vs. T2D (n=13) human donors. Data are normalized to *PC1* expression in healthy islets and *18S* was used as a housekeeping gene. **P<0.01.

### β cell-specific inactivation of furin in vivo

β cell-specific inactivation of furin (βfurKO) was achieved by intercrossing RIP-Cre^Herr^ mice lacking the human growth hormone (hGH) expression enhancer (21, 22), from here on referred to as RIP-Cre, with furin-floxed mice (23). In the latter mouse model, LoxP sites were introduced flanking exon 2, an essential exon encoding the translation initiation site, the signal peptide and part of the furin prodomain. We have previously shown that Cre-mediated excision of this sequence inactivates furin (23). The efficiency of this recombination event was evaluated by qRT-PCR on isolated islets. Using primers annealing to exon 2 and exon 3, only functional transcripts in which exon 2 was not excised were amplified. Islet furin expression was reduced by ~85% in βfurKO mice (**Figure S2A**). The remaining expression can be explained by detection of furin transcripts in the other cell types in the islets of Langerhans (mainly α and δ cells), in which Cre recombinase is not expressed (21). Since β cells in mice represent 60-80% of the cells in islets, these data suggest a complete or near-complete knockout of furin in β cells. There was no compensatory increase in mRNA expression of PCs that might provide redundancy for loss of furin activity (**Figure S2A**).

### βfurKO mice show elevated blood glucose levels and age-dependent glucose intolerance, without insulin resistance

Blood glucose levels were mildly but significantly elevated in 10-week-old βfurKO mice, both in fed (156 ± 5 mg/dL vs. 129 ± 3 mg/dL in controls, p<0.05) and 6h-fasted states (160 ± 5 mg/dL vs. 136 ± 5 mg/dL in controls, p<0.05) (**Figure S2B**). After overnight (16h) fasting, blood glucose levels were not significantly different. Intraperitoneal glucose tolerance tests (IPGTTs) were performed to assess glucose tolerance in male βfurKO and control (WT, Flox, Cre, heterozygous) mice **(Figure 2A-C)**. At the age of 10 weeks, βfurKO mice were significantly glucose intolerant, which worsened as the mice aged. Insulin tolerance tests at 10, 20 and 36 weeks of age did not show any significant differences, indicating normal peripheral insulin sensitivity **(Figure 2D-F)**. No significant differences in glucose tolerance were observed between βfurKO and control females at the age of 10 weeks, indicating that furin deficiency is not critical for glucose homeostasis in females at early age (**Figure S2C**). Glucose intolerance with normal insulin sensitivity was also observed when furin was knocked out using an islet-specific Ngn3-Cre driver line (**Figure S2D-E**). To quantify glucose stimulated insulin secretion (GSIS) *in vivo*, 24-week-old βfurKO and control male mice were injected intraperitoneally with a single glucose bolus (3 mg/g body weight). Analysis of plasma samples showed little or no increase in insulin plasma levels in βfurKO animals **(Figure 2G)**. Since glucose homeostasis was unaltered in Flox, Cre and heterozygous mice, potential Cre related effects of the RIP-Cre transgene and effects due to the introduction of *LoxP* sites in the *furin* gene can be ruled out. Therefore, only Flox controls were used throughout the rest of the study. It should be noted that Cre expression in this mouse model is not restricted to β cells, but also found in orexigenic RIP-expressing neurons in the hypothalamus (24). However, body weight was not significantly different between βfurKO and control mice on a normal chow diet (**Figure S2F**), suggesting that lack of furin does not have marked effects on body weight regulation by these neurons.

**Figure 2.**
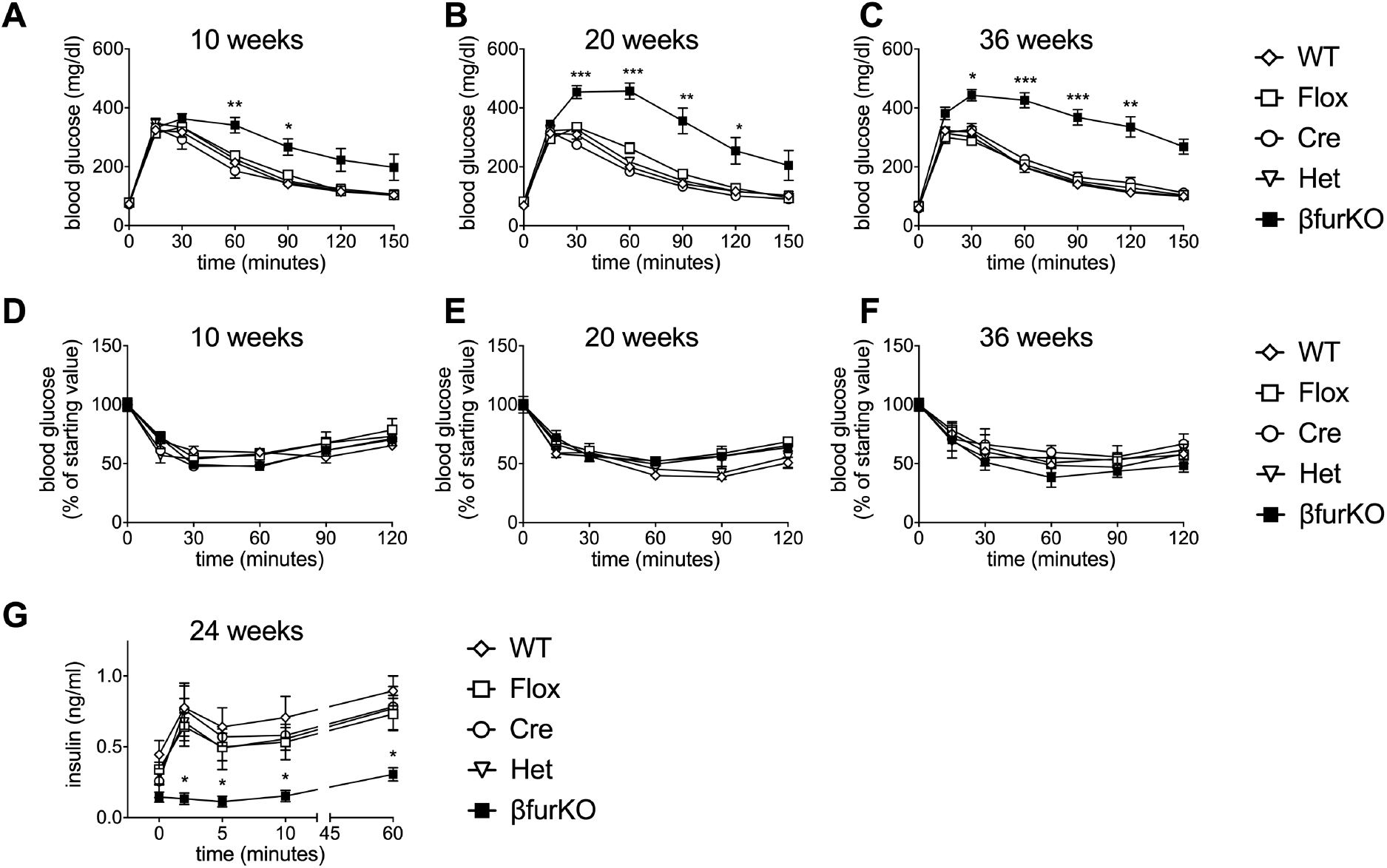
βfurKO mice show progressive glucose intolerance and impaired GSIS. (A-C) Intraperitoneal glucose tolerance tests (IPGTTs) on male βfurKO and control mice at 10 (A; n=6-17 mice/group), 20 (B; n=5-15 mice/group) and 36 (C; n=4-12 mice/group) weeks of age. (D-F) Intraperitoneal insulin tolerance tests (IPITTs) on βfurKO and control males at 10 (D), 20 (E) and 36 (F) weeks of age. (G) Twenty four-week-old βfurKO and control males were injected with a single bolus of 3mg/g BW D-glucose and insulin levels were measured at indicated time points, n=5-16 per group *P<0.05, **P<0.01, ***P<0.001, determined by 2-way ANOVA. All data are presented as mean ± SEM.

### Decreased β cell mass and islet insulin content in βfurKO mice

Islets from βfurKO mice appeared smaller than control islets during islet isolations. To substantiate this observation, β cell mass was quantified using an immuno-histological approach. β cell mass was significantly reduced in 24-week-old βfurKO mice (**Figure 3A**, 1.290 ± 0.303 mg vs. 2.572 ± 0.275 mg in controls, p=0.035). Similarly, β cell proliferation, measured as the percentage of Ki67^+^ β cells, was substantially reduced in βfurKO mice (**Figure 3B**, 0.13 ± 0.02 % for βfurKO islets versus 0.46 ± 0.02 % for control islets, p=0.0001). No differences in the percentage of apoptosis were detected by nucleosome ELISA **(Figure 3C)**. Insulin immunoreactivity was considerably weaker in βfurKO islets **(Figure 3D)** and decreased pancreatic and islet insulin levels were confirmed by ELISA **(Figure 3E-F)**.

**Figure 3.**
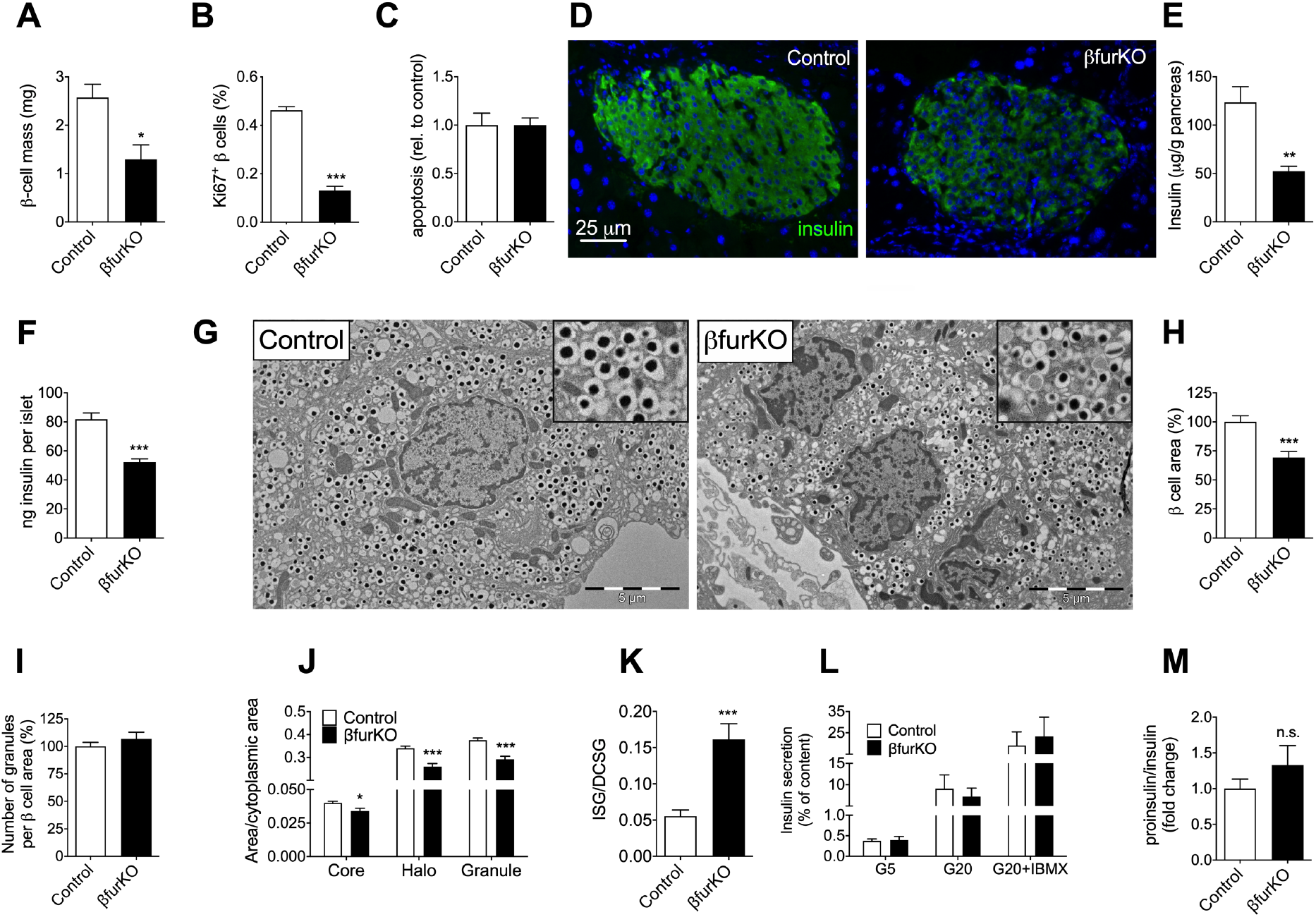
Decreased islet and pancreas insulin content, β cell mass, β cell proliferation, and ultrastructural abnormalities in 24-week-old βfurKO mice. (A) β cell mass quantification expressed in mg (n=3 mice/group). (B) Percentage of β cell proliferation as measured by Ki67 staining (n=3 mice/group). (C) β cell apoptosis as measured by nucleosome ELISA; data expressed as fold change compared to control values; n=4 mice per group. (D) Representative micrograph of insulin immunoreactivity in male βfurKO and control mice, scale bar = 25μm. (E) Pancreatic and (F) islet insulin content as measured by insulin ELISA; data are expressed as μg insulin per g pancreas (n=4-5 mice/group) and ng insulin per islet, respectively. (G) Representative electron micrographs of islets from 24-week-old control and βfurKO mice. Scale bar = 5 μm. (H) Quantification of total β cell area, (I) number of dense-core secretory granules per β cell cytoplasm, (J) β cell area occupied by dense-core secretory granules, and (K) immature/mature secretory granules (ISG/SG) ratio in control and furin knockout β cells (n=15 cells). (L) Islet GSIS, quantified as amount of insulin secreted in the medium corrected for total islet insulin content. G5, 5mM glucose; G20, 20mM glucose and G20 + IBMX. (M) Ratio of secreted proinsulin/insulin, n=4 mice/group; n.s., not significantly different. All data are represented as mean ± SEM, *P<0.05, **P<0.01, ***P<0.001.

Ultrastructural analyses by electron microscopy showed that furin-deficient β cells were smaller than controls **(Figure 3G-H)**. Despite similar numbers of secretory granules per cell area **(Figure 3I)**, granules in βfurKO islets were significantly smaller, showing reduced halo and core areas compared to controls **(Figure 3J)**. Furthermore, furin knockout β cells displayed a significant increase in the number of immature secretory granules (less electron dense, gray core) **(Figure 3K)**. These results suggest that furin is essential for β cell homeostasis and secretory granule biogenesis. There were no significant differences in GSIS of isolated islets when secreted insulin was corrected for total islet insulin content (**Figure 3L**). Similarly, the ratio (secreted proinsulin)/(secreted insulin) did not differ between genotypes (**Figure 3M**). These data suggest that there are no defects in insulin secretion or proinsulin processing in βfurKO islets, and instead reduced functional β cell mass is the main driver of the phenotype.

### Whole genome expression profiling and SILAC analysis show activation of an ATF4-dependent anabolic program in βfurKO islets

To explain the perturbations observed in this model, whole genome microarray analysis was performed on total islet RNA isolated from 12-week-old βfurKO and control mice. It is important to note that at this age βfurKO mice are glucose intolerant. Although this method does not directly provide furin targets, notion of differential gene expression might elucidate affected cellular processes and signaling pathways. Using this approach, downstream effects can be identified, potentially leading to upstream furin substrates.

Of the 28,853 genes examined with the GeneChip Mouse Gene 1.0 ST Array, 126 genes were differentially expressed in βfurKO islets with significance cut-off p<0.01 and fold change ≥1.5 (89 up, 37 down; **Figure 4A** and **Table S1**). Ingenuity Pathway Analysis (IPA, Qiagen) was performed to evaluate changes in biological processes (**Table S2**). In this context, mRNAs encoding AA transporters (*Slc1a4, Slc1a5, Slc35f2, Slc6a9, Slc7a3, Slc7a5, Slc7a11*), enzymes involved in AA metabolism (*Aass, Asns, Bcat1, Cth, Gpt2, Phgdh*) and aminoacyl-tRNA synthetases (*Eprs, Mars1*) were significantly upregulated **(Table S1)**.

**Figure 4.**
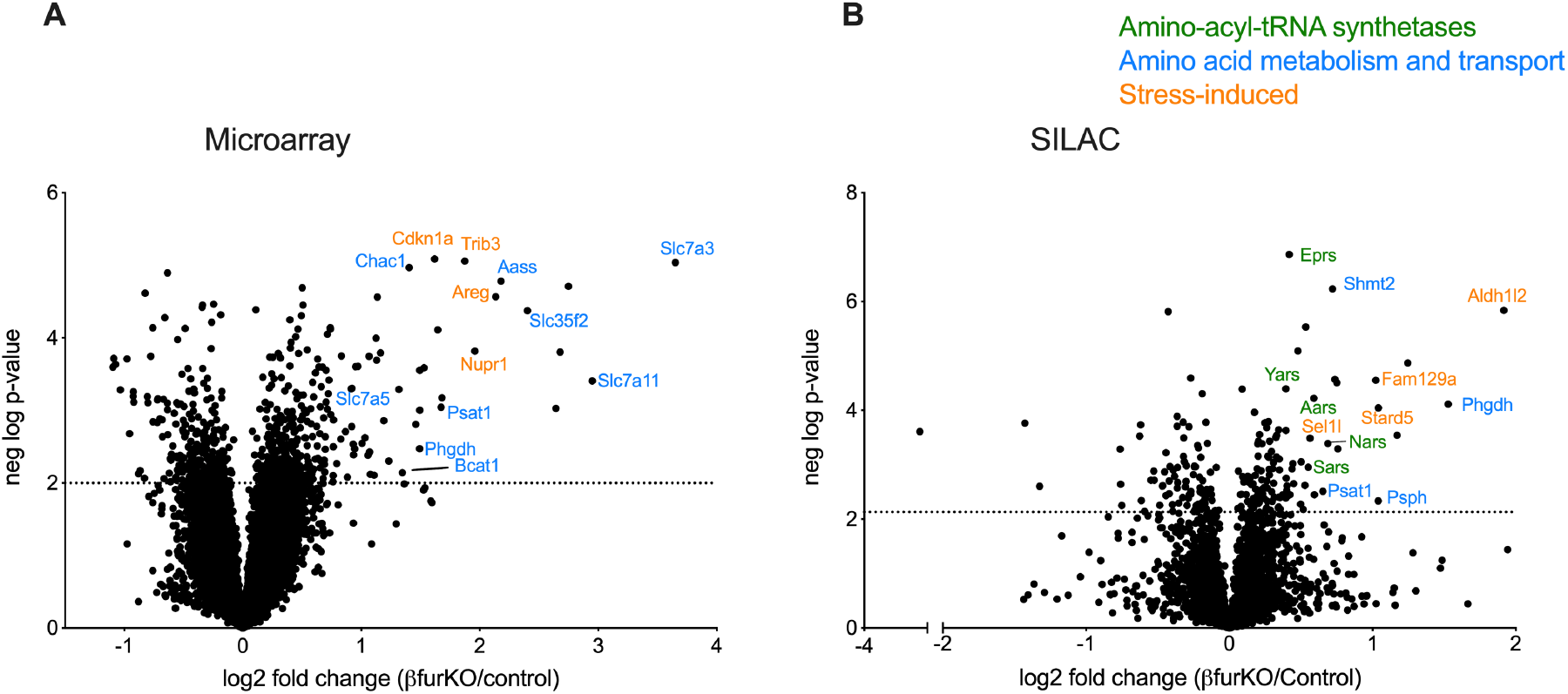
Whole-genome expression and quantitative proteomics analyses on βfurKO islets reveal a significant upregulation of ATF4 target genes. (A) Volcano plot showing differentially expressed genes in βfurKO vs. control islets (n=4 male mice per group). Negative log p-value = 2 was arbitrarily set as cut-off. (B) Volcano plot showing differentially expressed proteins in βfurKO vs. control islets.

To identify transcriptional regulators responsible for this observed gene expression profile, the IPA upstream regulator analysis tool was used. This tool predicted significant (p=2.86E-30) activation of Activating Transcription Factor 4 (ATF4) **(Table S3)**, a key transcription factor involved in the integrated stress response (25). It has previously been shown that ATF4 induces an anabolic transcription program in β cells during chronic ER stress, including expression of AA transporters and aminoacyl-tRNA synthetase genes (26). Consistently, other stress-induced genes (*Atf5*, *Chop* [=*Ddit3*], *Cdkn1a*, *Eif4ebp1*, *Fam129a*, *Nupr1*, *Sel1l*, *Trib3*, *Stard5*) were also upregulated (**Figure 4A** and **Table S1**).

To establish differences in the global protein profile, Stable Isotope Labeling of Amino acids in Cell culture (SILAC) analysis was performed on βfurKO and control islets using the murine *β* cell line Min6 as a spike-in standard. The minimum negative log p-value cutoff for statistical significance after multiple hypothesis correction was 2.13682 (p=0.0072). Using this condition, we found 203 differentially expressed proteins **(Figure 4B and Table S4)**. Consistent with our microarray data, IPA analysis showed significant changes in ‘metabolism of AA’ (p=7.19E-10) **(Table S5)**. In this context, AA synthesis enzymes (PHGDH, PSAT1, PSPH, SHMT2) showed increase in expression. In addition, IPA canonical pathway analysis showed activation of ‘tRNA charging’ (**Table S6**), with 9 aminoacyl-tRNA synthetase proteins in this pathway being significantly (p<0.0072) upregulated (AARS, CARS, EPRS, IARS, NARS, LARS, SARS, TARS, YARS) (**Figure 4B** and **Table S4**). Upstream analysis predicted activation of PKR-like ER kinase (PERK; EIF2AK3, p=3.21E-15; **Table S7**) and ATF4 (p=3.59E-11). PERK is an ER-proximal sensor of unfolded proteins that induces ATF4 translation via phosphorylation of Ser51 on eukaryotic initiation factor 2α (eIF2α). Together, these data suggest that β cells lacking furin induce an anabolic program involving upregulation of amino acid transporters and aminoacyl-tRNA synthetases, likely through activation of the ATF4 transcription factor. Consistent with data shown in **Figure 3F**, we observed a significant decrease in insulin I by means of SILAC (**Table S4**, ~30% decrease, p = 0.01). **Figure S3** shows a good correlation (r=0.4186, p<0.0001) of microarray and SILAC data for all proteins with at least a 25% increase or decrease in expression (n=73 proteins).

Negative log p-value was calculated to be 2.13682 after multiple hypothesis correction and was set as cut-off. Genes or proteins were color-coded according to their function.

### ATF4 upregulation in βfurKO islets is mediated by mTORC1 independently of eIF2α phosphorylation

Several kinds of stress conditions (ER stress, hypoxia and nutrient deprivation) increase phosphorylation of eIF2α, which leads to increased expression of ATF4 at the translational level and subsequent upregulation of ATF4 target genes. RT-qPCR expression analysis in islets confirmed increased expression of the ATF4 target genes *Chop* and *Trib3* (**Figure 5A**) whereas *Atf4* mRNA levels were unaltered. At the protein level, both ATF4 and CHOP were increased in βfurKO islets, confirming activation of ATF4 (**Figure 5B**). Surprisingly, βfurKO islets did not exhibit increased phosphorylation of PERK or eIF2α (**Figure 5B**) nor were other ER stress markers increased (**Figure S4A-B**). Emerging evidence suggests that ATF4 can be activated by mammalian target of rapamycin complex 1 (mTORC1) **(Figure 5C)** (27–29). In agreement with this idea, phosphorylation levels of p70 ribosomal protein S6 kinase (S6K) and eukaryotic translation initiation factor 4E-binding protein 1 (4E-BP1), two substrates of mTORC1, were augmented in βfurKO islets **(Figure 5D)**. This suggests an activated mTORC1-ATF4 axis in islets from βfurKO mice.

**Figure 5.**
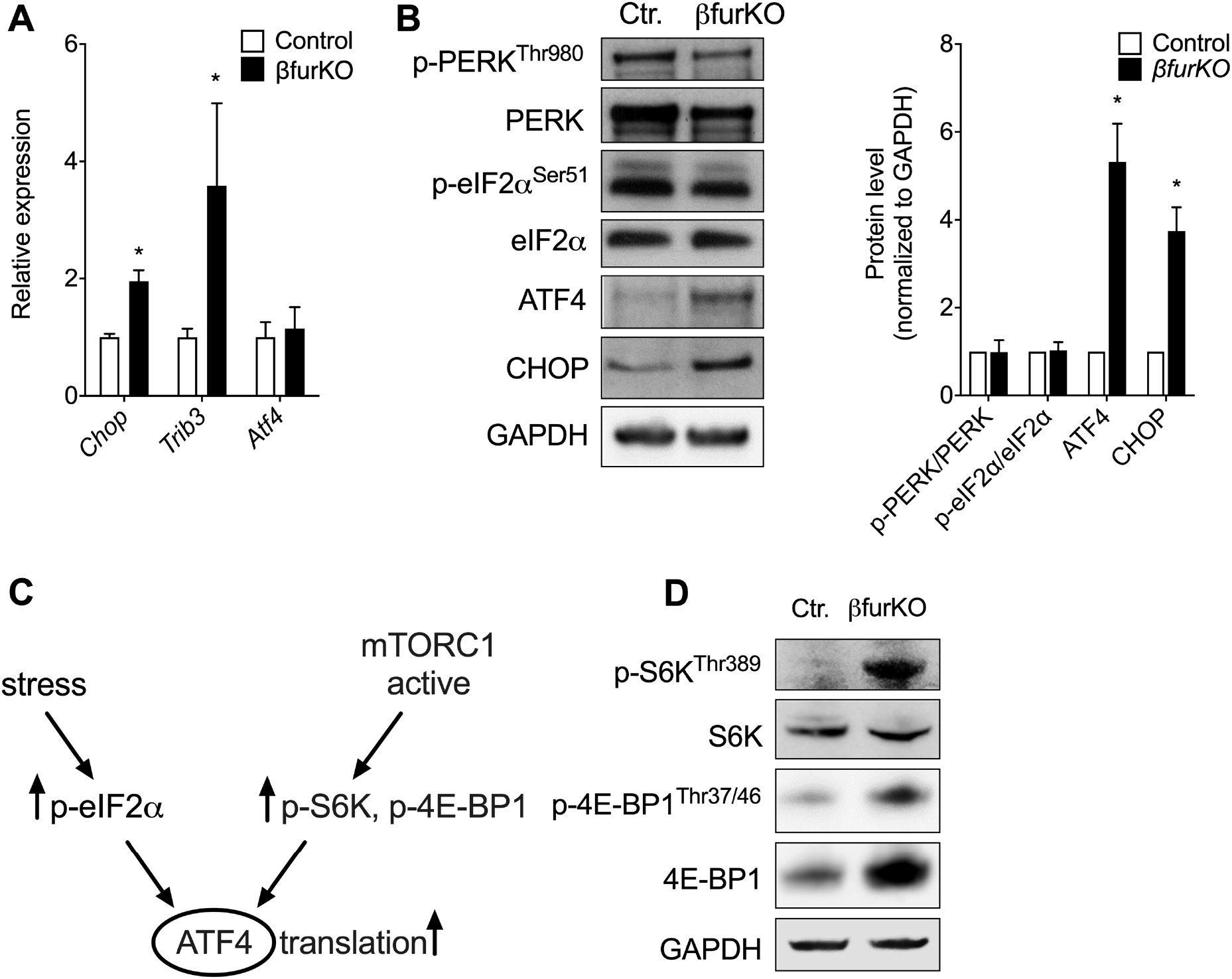
Furin knockout islets exhibit activation of the ATF4 pathway and mTORC1 activity, but no increase in ER stress markers. (A) Quantitative RT-PCR analyses of the mRNA levels of *Atf4* and target genes *Chop* and *Trib3* in whole islet lysates from control and βfurKO mice, n=3-7 per group. Values were normalized to *Gapdh* expression. (B) Immunoblot analyses and quantification of p-PERK, total PERK, p-eIF2α, total eIF2α, ATF4 and CHOP protein levels in whole islet lysates. P-PERK/PERK, p-eIF2α/eIF2α, ATF4 and CHOP signals were normalized to GAPDH. Data represent the mean of 3 independent experiments. (C) Mechanisms that increase ATF4 translation. (D) Immunoblot analyses of p-S6K and p-4E-BP1 protein levels in whole islet lysates from control and βfurKO mice. For immunoblot analyses GAPDH was used as the loading control. Data are represented as mean ± SEM, *P<0.05, **P<0.01, ***P<0.001.

### Furin knockout βTC3 cells exhibit reduced cell proliferation and upregulation of the ATF4 pathway

To substantiate our *in vivo* findings in a cellular model, we generated a furin knockout β cell line lacking furin using the CRISPR-Cas9 system **(Figure 6A)**. Cell proliferation was significantly reduced in furin knockout βTC3 cells **(Figure 6B)**. In addition, furin knockout βTC3 cells showed increased *Trib3* mRNA **(Figure 6C)** and CHOP and ATF4 protein levels **(Figure 6D)**, suggesting ATF4 activation. Like βfurKO islets, we did not observe increased phosphorylation of eIF2α compared to control cells in furin knockout βTC3 cells **(Figure 6D)**, implying that ATF4 upregulation is not mediated by eIF2α in this cell line. Additionally, furin knockout βTC3 cells displayed higher mTORC1 activity as demonstrated by the increased phosphorylation levels of S6K and 4E-BP1 **(Figure 6E)**. Together, we conclude that these furin knockout βTC3 cells can be used as a representative model to investigate the molecular mechanisms caused by furin deficiency in β cells *in vivo*. To substantiate that activation of mTORC1 causes the ATF4 upregulation in the furin knockout βTC3 cell line, we performed a rescue experiment with rapamycin, which specifically inhibits mTORC1 at short incubations. After treatment with rapamycin, ATF4 protein levels were reduced compared to control βTC3 cells **(Figure 6F-G)**.

**Figure 6.**
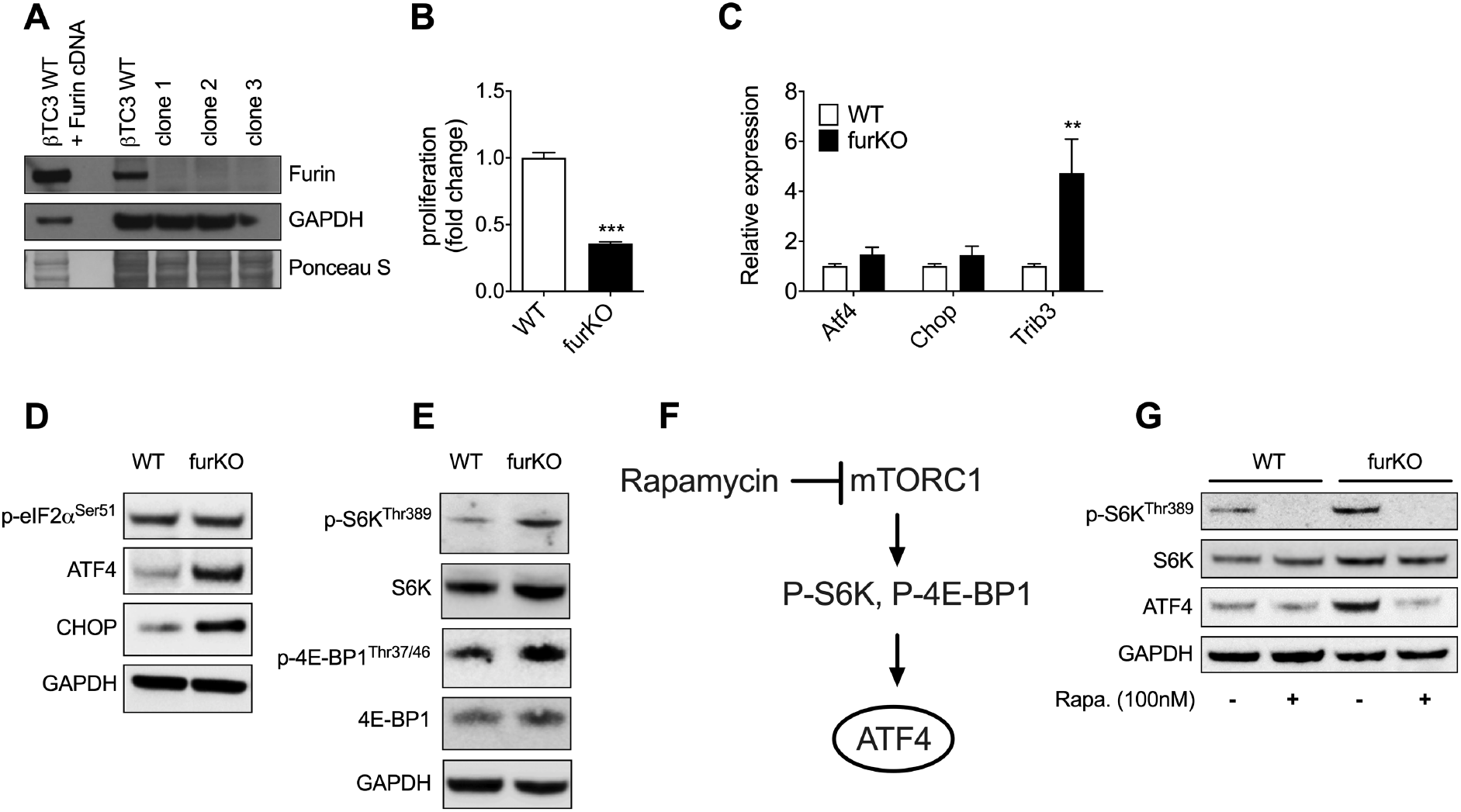
Furin knockout βTC3 cells exhibit activation of ATF4 and downstream genes in an mTORC1-dependent but PERK/eIF2α-independent manner. (A) Protein immunoblotting of cell extracts from control and furin knockout βTC3 cells. GAPDH and Ponceau S staining were used as loading control. (B) Cell proliferation measured with a WST-1 colorimetric assay in control and furin knockout βTC3 cells. Data, expressed as fold change over control cells, are the mean ± SEM of three independent experiments (n=9) using three different clones. (C) Quantitative RT-PCR analyses of the mRNA levels of *Atf4*, *Chop* and *Trib3* in whole βTC3 lysates. (D) Immunoblot analyses of p-eIF2α, ATF4 and CHOP protein levels in whole βTC3 lysates. (E) Immunoblot analyses of p-S6K, total S6K, p-4E-BP1, total 4E-BP1 protein levels in whole βTC3 lysates. (F) Blocking the mTORC1 pathway by rapamycin is expected to inhibit downstream substrates and to decrease translation of ATF4. (G) Immunoblot analyses of p-S6K, total S6K, and ATF4 protein levels in whole cell lysates from βTC3 cells treated either with vehicle (0.01% DMSO) or 100 nM rapamycin for 16 h. For immunoblot analyses GAPDH was used as the loading control and results represent three independent experiments. For quantitative RT-PCR analyses the values were normalized to *Gapdh* expression and the data are represented as mean ± SEM, n=3-6 per group, *P<0.05, **P<0.01, ***P<0.001.

### The V-ATPase and mTORC1 signaling: involvement of Ac45 and PRR

We next sought to determine the mechanism driving mTORC1 hyperactivation in βfurKO cells. We first focused on the V-ATPase proton pump, as it is known to influence mTORC1 signaling in different ways. Recent data have shown that it serves as part of a docking and activation site for mTORC1 on lysosomes (30). In addition, dysfunction of lysosomal acidification results in impaired lysosomal degradation and overactivation of mTORC1 signaling in a mouse model of amyotrophic lateral sclerosis (ALS)/frontotemporal dementia (FTD) (31). Two critical V-ATPase accessory subunits, ATP6AP1/Ac45 and ATP6AP2/(Pro)-renin receptor (PRR) are known furin substrates (17, 32) and highly expressed in β cells in mice (33) and humans (20). As such, we hypothesized that impaired Ac45 and/or PRR cleavage would lead to disturbed lysosomal acidification hereby affecting mTORC1 activity in β cells. We first investigated the role of the V-ATPase in mTORC1 signaling by treating wild type βTC3 cells with the V-ATPase inhibitor Bafilomycin A1 for 24h. This resulted in a phenocopy of the βfurKO model showing activation of mTORC1 pathway and the upregulation of ATF4 **(Figure 7A-B)**. Next, we investigated the processing of Ac45 and PRR in furKO cells. The furin cleavage sites of the corresponding precursors were shown to be at the RVAR^231^ and RKSR^278^ sites respectively (17, 32) (**Figure 7C**), while PRR can also be cleaved by ADAM19 (34), and by SKI-1 at a RTIL^281^ site downstream of the furin cleavage site (35). Mature Ac45 is heavily glycosylated and appears as a broad smear around 45 kDa, but the cleaved form can be detected as a 24-kDa peptide when treated with N glycosidase F (17) (**Figure 7D**, left panel). Using this treatment, we show a complete lack of the 24-kDa form, indicating that Ac45 cleavage does not occur in furKO β cells (**Figure 7D**, right panel). The intracellular amount of cleaved PRR did not appear to be affected (**Figure 7E,** upper panel), but the soluble secreted form (sPRR) was significantly reduced in the medium of furKO β cells (**Figure 7E**, lower panel). Knockdown of *Atp6ap1*/*Ac45* showed a non-significant trend towards increased phosphorylation levels of S6K and 4E-BP1 and upregulation of ATF4. No indications for activation of the mTORC1 pathway were observed after knockdown of *Atp6ap2/Prr*. (**Figure 7F-G)**. Likely, these results reflect the compound nature of the phenotype, whereby loss of furin results in impaired cleavage for several substrates, leading to the phenotype. On the other hand, the uncleaved proprotein might lead to additional effects that a genetic knockdown cannot replicate.

**Figure 7.**
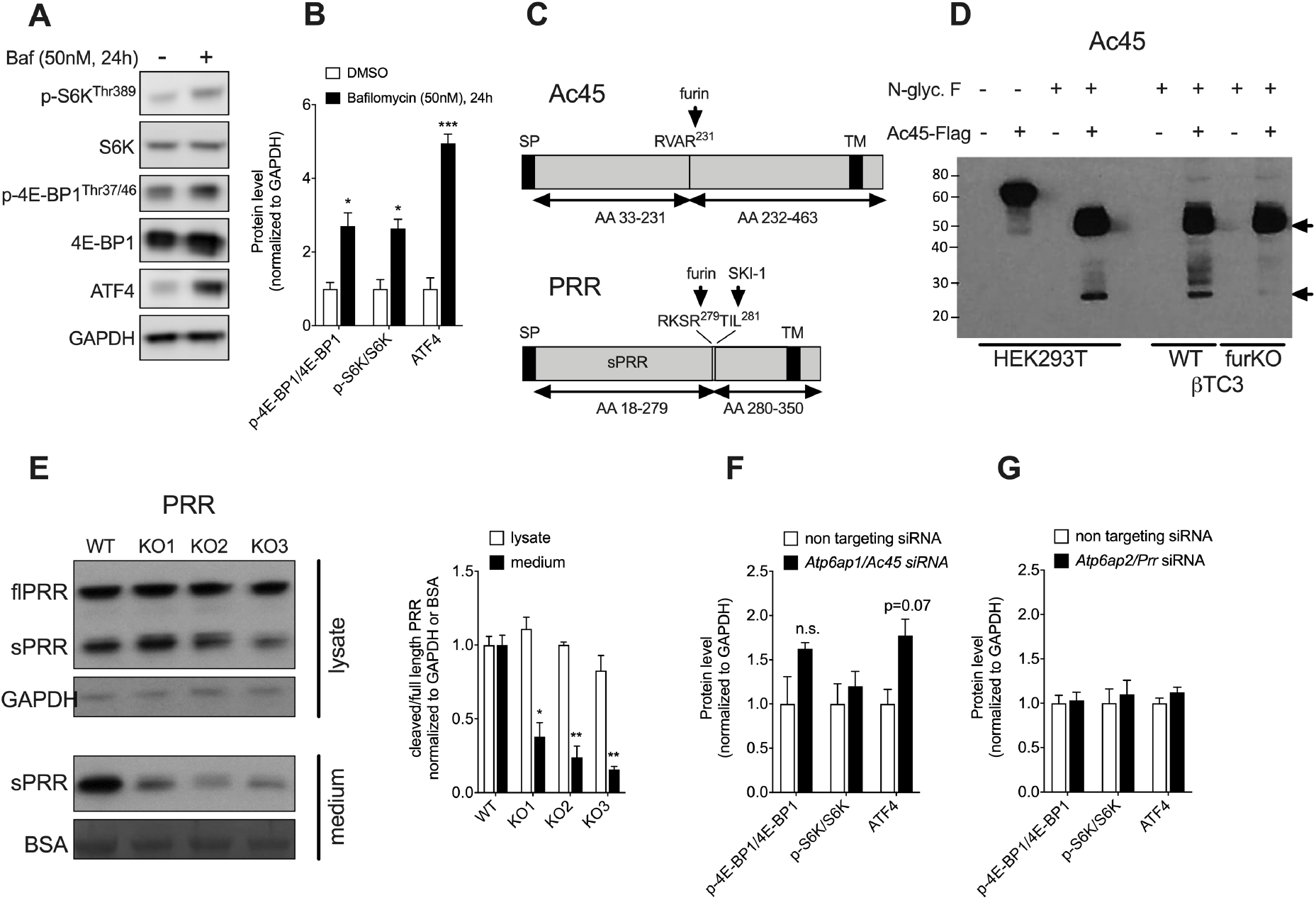
Effect of V-ATPase inhibition on mTORC1/ATF4 pathway in βTC3 cells and processing of Ac45 and PRR in βTC3 (WT and furKO) cells. (A) Western blot for p-S6K, total S6K, p-4E-BP1, total 4E-BP1, ATF4 in βTC3 cells incubated with the V-ATPase inhibitor bafilomycin A1 (50 nM final concentration) for 24h. GAPDH was used as a loading control. (B) Western blot quantification from n=3 independent experiments. (C) Schematic illustration of Ac45 and PRR protein structures. Furin and SKI-1 cleavage sites are indicated by arrows. SP, signal peptide; TM, transmembrane domain; AA, amino acids, SKI-1 (Subtilisin Kexin Isozyme-1). (D) HEK293T and βTC3 (WT and furKO) cells were transfected with a Flag-tagged Ac45 construct. Western blot of HEK293T cell lysate before and after N-glycosidase F (N-glyc F) treatment, with the processed form (24 kDa) appearing only after deglycosylation (left HEK293T panel). Western blot of βTC3 cell lysate, wild type (WT) and furin knockout (furKO), both treated with N-glyc F (right panel). The indicated positions of proAc45 and Ac45 correspond to the predicted MW of the deglycosylated peptide backbone (46 kDa and 24 kDa respectively, black arrows). (E) Full length (fl) and soluble (s)PRR levels in the lysate (upper panel) and conditioned medium (lower panel) of furin knockout (furKO) and WT βTC3 cells. Three independent furKO clones were assessed. Quantifications on the right-hand side show n=3 independent experiments. *P<0.05, **P<0.01. (F) Western blot quantification of p-4E-BP1/4E-BP1, p-S6K/S6K and ATF4 in βTC3 cells that were subjected to *Atp6ap1*/*Ac45* knockdown by siRNA, n=4 independent experiments, signals were normalized to GAPDH. (G) Western blot quantification of p-4E-BP1/4E-BP1, p-S6K/S6K and ATF4 in βTC3 cells that were subjected to *Atp6ap2*/*Prr* knockdown by siRNA, n=4 independent experiments, signals were normalized to GAPDH. The results are represented as mean ± SEM *P<0.05, **P<0.01, ***P<0.001.

### Lack of processing or absence of the insulin receptor does not lead to activation of the mTORC1/ATF4 axis

Upstream regulator analysis of the SILAC dataset by IPA predicted significant (P=0.0005) inhibition of the insulin receptor (IR) (**Table S6**), a potential PC substrate (23, 36). The IR can be proteolytically processed by furin at an RKRR site, yielding mature α and β subunits (**Figure 8A**), and this cleavage event is necessary for insulin binding, internalization and autophosphorylation of the receptor (37, 38). Moreover, insulin signaling activates mTORC1 by effectors downstream of the IR (39). The previously reported β cell specific IR knockout mouse model (βIRKO) displays a phenotype similar to βfurKO mice with progressive glucose intolerance and smaller islets, and increased pS6K in islets (40, 41). We therefore hypothesized that uncleaved IR might lead to altered mTORC1 activity. The absence of furin in the βTC3 cell line resulted in the lack of IR processing (**Figure 8B**) and transfection with recombinant furin rescued the cleavage of mature IR (**Figure 8C**). Furthermore, we found that furin knockout βTC3 cells showed little response to insulin stimulation (**Figure 8D**). To test the hypothesis that the activation of the mTORC1-ATF4 pathway in furin knockout β cells is caused by the lack of IR cleavage, we intercrossed RIP-Cre mice (21) with IR-floxed animals, resulting in βIRKO animals, which showed a ~68% reduction of *Insr* mRNA in islets (**Figure 8F**). In contrast to the earlier observation in βIRKO mice generated using the RIP-Cre^Mgn^ driver (40), glucose tolerance was only mildly affected and only in 24-week-old animals (**Figure 8E**). This is considerable milder than βfurKO mice from a similar age (20 weeks; **Figure 2B**). Genes downstream of ATF4, *Trib3* and *Chop*, were not induced in βIRKO islets and even showed a trend towards reduced expression (**Figure 8F**). Knockdown of *Insr* in βTC3 cells did not activate the mTORC1 pathway either (**Figure 8G**). These results show that lack of IR and downstream signaling does not lead to an induction of mTORC1 activity and downstream activation of ATF4 in β cells.

**Figure 8.**
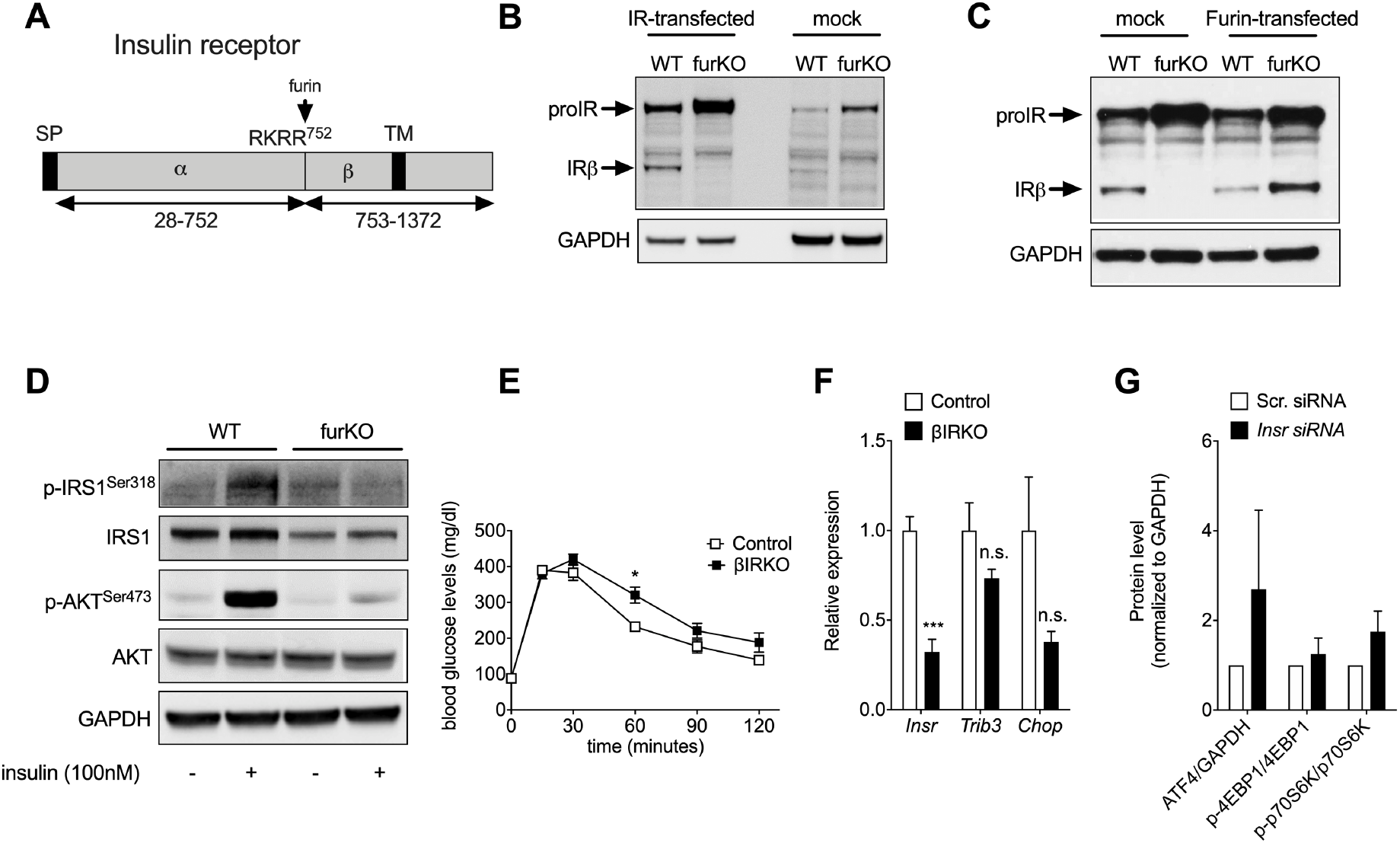
Lack of insulin receptor processing and impaired insulin signaling in furin-deficient β cells. (A) Schematic of insulin receptor structure. The pro-receptor undergoes furin cleavage at the indicated tetrabasic site. SP, signal peptide; α, alpha subunit; β, beta subunit; TM, transmembrane domain, AA, amino acids. (B) Immunoblot analyses of IR protein levels in whole cell lysates from IR-transfected or non-transfected βTC3 cells. (C) Immunoblot analyses of IR protein levels in whole cell lysates from βTC3 cells transfected with IR or co-transfected with IR and furin. (D) Immunoblot and quantification of the phosphorylation and abundance of IRS1 and Akt in whole cell lysates from βTC3 cells treated with 100 nM insulin or vehicle for 5 min. (E) IPGTT of βIRKO versus control (Flox) males, 24 weeks old, n=8-16. (F) Islet qRT-PCR of βIRKO, for *IR*, *Trib3* and *Chop* genes, n=4/group. (G) Western blot quantification of ATF4/GAPDH, p-4E-BP1/total 4E-BP1 and p-S6K/total p-S6K protein levels in whole cell lysates from βTC3 cells transfected with either scramble siRNA or mouse IR siRNA. GAPDH was used as the loading control and results represent 3-4 independent experiments, reported as mean ± SEM. *P<0.05, **P<0.01, ***P<0.001.

## Discussion

In this study we have demonstrated that furin is essential for β cell function, and that the dysregulation of its activity can affect β cells through the induction of the stress factor ATF4 in a mTORC1-dependent manner (**Figure 9**). Primary phenotypic observations in βfurKO mice include age-dependent, progressively deteriorating glucose tolerance, with mild hyperglycemia in fed and 6-hour fasted states. βfurKO mice also show defective first phase GSIS, a trait that is observed in patients with impaired glucose tolerance and in early stages of T2D (42). Since we observed β cell mass reduction and diminished insulin content in βfurKO mice, the number of β cells might be below the threshold required to maintain adequate glucose homeostasis, which could directly lead to impaired pulsatile insulin secretion (42, 43).

**Figure 9.**
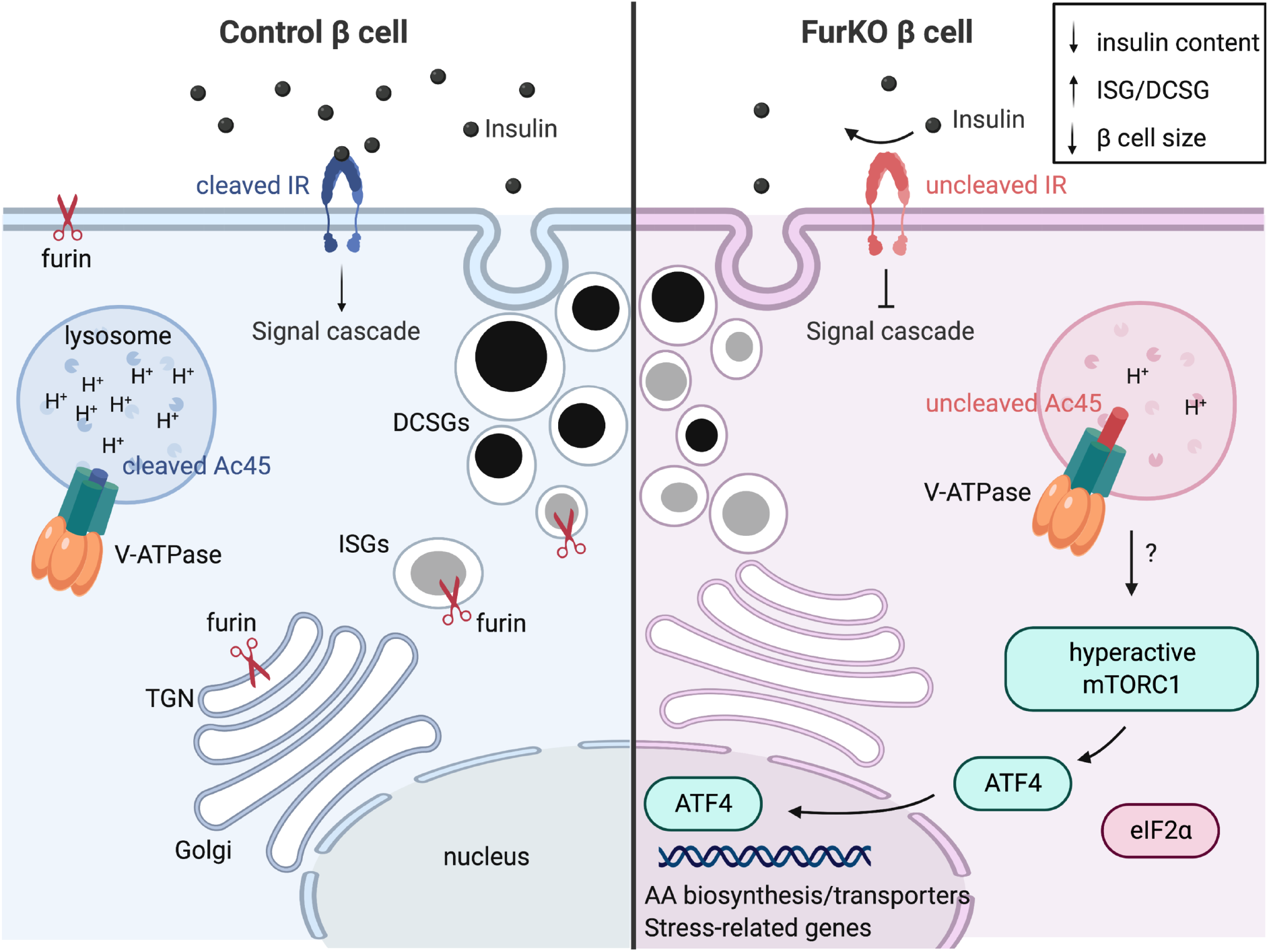
Schematic overview of phenotypes in β cells lacking furin. Furin is a proprotein convertase concentrated in the *trans*-Golgi network (TGN) and cycles through a complex trafficking circuit that involves several TGN/endosomal compartments and the cell surface. In (neuro)endocrine cells, furin is also present in insulin-containing immature secretion granules (ISGs) and is removed and returned to the TGN before ISGs mature to dense-core secretion granules (DCSGs). β cells lacking furin (furKO) show reduced cleavage of Ac45 - an essential subunit of the V-ATPase proton pump - and insulin receptor (IR), resulting in reduced acidification of intracellular organelles (e.g. lysosomes) and blunted insulin signaling, respectively. FurKO cells induce an expression program that involves amino acid (AA) biosynthesis enzymes and transporters, and stress-related genes. This program is induced by a mammalian target of rapamycin complex 1 (mTORC1) - Activating Transcription Factor 4 (ATF4) axis, in an eIF2α independent manner. FurKO cells show increased ISG/DCSG ratio, are smaller and exhibit reduced insulin content, resulting in impaired glucose tolerance in the whole organism.

Whole-genome expression profiling and proteomics analyses pointed towards activation of the transcription factor ATF4. Translation of this transcription factor can be induced by ER stress, via the PERK-eIF2α-ATF4 pathway (44). However, we did not detect activation of this pathway, nor of the other two signaling arms of UPR, the IRE1α-XBP1 and the ATF6 pathways. Instead, activation of mTORC1 was identified as upstream activator of ATF4 in βfurKO islets. Consistent with our data, studies in other cell types have shown that ATF4 can be activated by mTORC1 independently of its canonical induction via phosphorylated eIF2α. For instance, the proteasome inhibitor bortezomib promotes the transcription and translation of ATF4 in an mTORC1-dependent manner (28). Moreover, ATF4 stimulates the *de novo* purine synthesis pathway in an mTORC1-dependent and eIF2α-independent manner in various mouse and human cell lines (27). Interestingly, Park and colleagues have shown that mTORC1 transcriptionally regulates amino acid transporters, metabolic enzymes and aminoacyl-tRNA synthetases through post-transcriptional control of ATF4, which was independent of eIF2α phosphorylation (29). Although the evidence of mTORC1-ATF4 axis has recently been documented, the importance of this pathway in β cells and more specifically in T2D has not been explored yet.

In pancreatic β cells, mTORC1 plays major roles in β cell function and mainly participates in β cell proliferation and β cell mass regulation (45). However, whereas at early stage mTORC1 activation induces β cell hypertrophy resulting in hypoglycemia and hyperinsulinemia, chronic hyperactivation of mTORC1 in β cells promotes progressive hyperglycemia and hypoinsulinemia accompanied by a reduction in β cell mass (46). Moreover, chronic hyperactivation of mTORC1 promotes ER stress and causes impaired autophagic response (47), leading to β cell failure. Therefore, it is highly likely that mTORC1 hyperactivity leads to β cell dysfunction in βfurKO mice. Human islets isolated from T2D patients exhibit increased mTORC1 activation (48). Genetic or chemical inhibition of mTORC1-S6K signaling restores insulin secretion in human islets from patients with T2D, suggesting that hyperactive mTORC1 impairs β cell function (48). As such, our βfurKO mouse could serve as a new genetic model to study mTORC1 hyperactivity in β cells.

Loss of furin activity in β cells potentially affects cleavage of several substrates that contribute to the observed phenotype to different extents. Using a data driven approach, we investigated candidate substrates based on a potential link with the mTORC1-ATF4 axis. Proteins involved in V-ATPase function are likely candidates, since the V-ATPase complex acts as an mTORC1 activation site (30, 49). Furin activity has already been shown to be critical for V-ATPase function and acidification in mice (17) and in yeast (50). Recently, a cryo-EM model of the rat V-ATPase complex was published, showing Ac45 and PRR present as cleaved forms interacting with subunits that line the proton permeable pore (51). In addition, mass spectrometry analyses on V-ATPase preparations showed that predominantly the cleaved forms of Ac45 and PRR are present (51). Here, we show almost complete lack of Ac45 cleavage in furin deficient β cells. As an essential subunit regulating V-ATPase function (17, 52), Ac45 might modify β cell mTORC1 activity by relaying amino acid sufficiency from the lysosomal lumen. In patients with follicular lymphoma, mutations that activate mTORC1 often co-occur with mutations in *ATP6AP1/AC45* (53). It was hypothesized that some of these *ATP6AP1/AC45* mutants might convey a false amino acid sufficiency signal or alter interactions between V-ATPase and components of the mTORC1 complex. As such, lack of Ac45 cleavage in β cells might explain why we observe mTORC1 hyperactivity. In addition, accumulating evidence shows the importance of the prorenin receptor (PRR) in pancreatic β cells. Similar to Ac45, PRR is an essential V-ATPase subunit and highly expressed in this cell type (54). Interestingly, it was identified as an interactor for the glucagon-like receptor 1 (GLP1) in human and mouse islet screens (54). Similar to furin and Ac45 (17, 52), knockdown of *Atp6ap2/Prr* in β cells results in reduced granular acidification (54). Mice lacking PRR in β cells exhibit altered glucose tolerance, reduced insulin content and show accumulation of very large multigranular vacuoles in the cytoplasm (33). While we did not observe these structures in βfurKO islets, it does not rule out a contribution of PRR to the phenotype. β cells lacking furin exhibit reduced shedding of sPRR. The exact timing and mechanism of cleavage is still a matter of debate and might be cell type dependent (32, 34, 35). Whether the reduced amount is the consequence of reduced cleavage or reflects reduced protein stability of the soluble form remains to be established. Finally, we did not observe significant upregulation of mTORC1 activity upon Ac45 or PRR knockdown. This finding does not necessarily rule out Ac45 and PRR as candidate substrates and might instead reflect the fact that the efficiency of gene knockdown is insufficient to obtain a phenotype or that the cause if multifactorial. The phenocopy obtained with bafilomycin suggests a crucial role for the V-ATPase.

The previously published β cell-specific IR knockout model (βIRKO) shows reduced GSIS *in vivo*, a progressive impairment in glucose tolerance, reduced pancreatic insulin content and reduced β cell proliferation (40), phenotypes that are also observed in βfurKO mice. However, these studies made use of RIP-Cre^Mgn^ driver line, which was later shown to exhibit glucose intolerance, decreased insulin secretion and β cell hypoplasia in younger mice and hyperplasia in older mice (55, 56), mediated by ectopic hGH expression (22). In addition, Cre expression in RIP-Cre^Mgn^ mice is not restricted to β cells and is also found to be widespread in the brain (57), which likely contributes to the phenotypes as well. Using the RIP-Cre^Herr^ driver line, we observed a phenotype that was considerably milder compared to these previous studies (40). In recent work using an inducible MIP1-Cre knock-in mouse line, inactivation of the IR in adult β cells resulted in normal glucose tolerance, insulin release and β cell mass when fed a normal chow diet, and became glucose intolerant only when challenged with a high fat diet (58). The current study extends that observation to genetic inactivation of the IR starting during development. Together, it shows that the earlier findings were likely confounded by the Cre driver line. As furin activity is necessary for IR function, and insulin signaling induces mTORC1, one would expect decreased mTORC1 activity in β cells lacking furin. However, impaired V-ATPase function might be dominant or uncleaved insulin receptor might convey different signals eventually leading to mTORC1 activation in β cells. Finally, we did not test the efficiency of Cre-mediated furin recombination in RIPHER neurons, so we cannot entirely rule out any phenotypic contribution by hypothalamus-specific furin inactivation. However, the phenocopy observed in a β cell line generated by CRISPR/Cas9 editing strongly suggests a defect at the level of the β cell.

Besides reduced cleavage of several substrates which contribute to the phenotype, the absence of furin itself may contribute as well. Furin is sorted to immature secretory granules (ISGs) but is excluded from mature secretory granules (59). Retrieval from ISGs requires AP-1 adapter complex and clathrin. The absence of furin as receptor for retrieval of soluble content from ISG during maturation might contribute to the increased number of ISG and morphological changes in the mature granules (60).

In conclusion, we describe the importance of the proprotein convertase furin in the regulation of β cell mass and function *in vivo*, likely related to its function in regulating the V-ATPase activity. Furthermore, we report that lack of furin results in activation of the stress-induced transcription factor ATF4 mediated by mTORC1. As such, this study sheds light on a novel molecular mechanism for the regulation of β cell function by furin and provides evidence that it is indeed the prime candidate in the PRC1 susceptibility locus for T2D.

## Experimental procedures

### Human islets

Human islets from healthy (n=10; HbA1c < 6,0) and T2D donors (n=13; HbA1c > 6,5 at diagnosis) were obtained from the Alberta Diabetes Institute Islet Core (University of Alberta). **Table S8** contains general information about the donors and additional information is available at https://www.epicore.ualberta.ca/IsletCore/. Islets were cultured overnight until they were handpicked and snap frozen. Protocols were approved by the Human Ethics Committee of IRCM.

### Mice

RIP-Cre^+/−^ mice were kindly donated by Dr. Pedro L. Herrera (University of Geneva Medical School) (21). These mice were bred with animals in which the essential exon 2 of the *furin* gene is flanked by loxP sites (23). This generates wild-type (‘WT’, RIP-Cre^−/−^;furin^wt/wt^), floxed (‘Flox’, RIP-Cre^−/−^;furin^flox/flox^), Cre control (‘Cre’, RIP-Cre^+/−^;furin^wt/wt^), β cell-specific furin heterozygous (‘Het’, RIP-Cre^+/−^;furin^flox/wt^) and β cell-specific furin knockout (‘βfurKO’, RIP-Cre^+/−^;furin^flox/flox^) mice. Insulin receptor floxed mice (IR^flox/flox^) were kindly provided by Dr. Jens Brüning (Max Planck Institute for Metabolism Research, Germany) and were intercrossed with RIP-Cre^+/−^ mice to obtain RIP-Cre^+/−^;IR^flox/flox^ animals. Mice were backcrossed at least 8 times to a C57Bl6 background (Janvier). Only male mice were used for experiments throughout the study. Mice were sacrificed using cervical dislocation. All experiments were approved by the KU Leuven Animal Welfare Committee, following the guidelines provided in the Declaration of Helsinki (KU Leuven project number 036/2015).

### Intraperitoneal glucose (IPGTT), insulin (IPITT) tolerance tests and glucose stimulated insulin secretion (GSIS)

Male WT, Flox, Cre, Het and βfurKO mice fed a normal chow diet were fasted overnight and injected with 2 mg/g body weight (BW) D-glucose (IPGTT) or 0.75 IU/g BW human insulin (ITT) in PBS, respectively. Blood glucose levels were monitored at indicated time-points using a Contour Glucometer (Roche). Mice were analyzed at ages 10 weeks (n=6-17 per group), 20 weeks (n=5-15 per group) and 36 weeks (n=4-12 per group) as indicated. In a separate experiment, 24-week-old mice (n=6-16) were fasted overnight and injected with a single intraperitoneal bolus of 3 mg/g D-glucose dissolved in PBS. Subsequently, plasma samples were collected at indicated time points and analyzed for insulin using the Ultrasensitive Mouse Insulin ELISA (Mercodia).

### Islet isolation

Islets were isolated by locally injecting the pancreas with 1 Wünsch unit/ml Liberase (Roche) in HANKS buffer as previously described (22). Injected pancreata were incubated for 18 min at 37°C in a shaker (220 rpm) and subsequently the islet fraction was separated from exocrine tissue using a Dextran T70 gradient. Finally, islets were handpicked twice in HANKS buffer under a stereomicroscope to reach a pure islet fraction for further processing.

### Islet Insulin Content and Release

For insulin secretion measurements, size-matched islets (n = 5 islets per tube, in triplicate for each condition per mouse, n=4 mice per genotype) were placed in glass tubes containing HEPES Krebs solution 0.5% BSA supplemented with glucose: 5 mM (G5), 20 mM (G20), or G20 with 250 mM IBMX (Sigma). Supernatant was collected after 1 h incubation at 37°C. The islets were sonicated for 3 min after adding acid ethanol (final concentration: 75% EtOH, 0.1 N HCl, 1% Triton). Samples were stored at 20°C until quantification. The ELISA kit used for insulin determination was from Crystal Chem. The ELISA kit used to quantify proinsulin was from Mercodia.

### β cell mass quantification

β cell mass of 24-week-old male mice was determined using a previously described protocol (22). Pancreatic sections sampled every 200 μm were stained for insulin using standard IHC protocols. Briefly, sections were heated for 20 min in Target Retrieval Solution (Dako), blocked in 20% fetal calf serum (FCS) and incubated with anti-insulin (1/10, Dako) in Antibody Diluent for 2h, followed by peroxidase-coupled secondary antibodies for 1h. 3’-3-Diamino-benzidine (DAB+, Dako) was used as substrate chromogen, after which sections were counterstained with hematoxylin and mounted. Six insulin-stained sections per mouse were photographed using a Zeiss Axioimager (Zeiss). Axiovision software (Micro Imaging, Heidelberg, Germany) was used to determine the relative insulin-positive area for every section. Subsequently, the β cell mass (mg) was calculated by multiplying relative insulin-positive area by the weight of the pancreas in mg.

### Immunofluorescence (IF)

Pancreata were isolated and fixed in 4% paraformaldehyde (PFA) in PBS at 4°C overnight. Subsequently, they were dehydrated by a graded series of ethanol and butanol and embedded in paraffin. Sections were cut at 5μm thickness. Sections were deparaffinized, rehydrated and boiled for 20 minutes in Target Retrieval solution (Dako) to recover epitopes. Slides were blocked with 20% FCS in PBS and incubated overnight with 1/10 anti-insulin (Dako) and 1/500 anti-Ki67 (Abcam) in Antibody Diluent (Dako) and subsequently incubated with Alexa antibodies (1/500) for 1h at room temperature.

### Pancreatic insulin content

Total pancreatic insulin content was determined using the acid/ethanol extraction method. Briefly, pancreata were homogenized in acid/ethanol (0.12 M HCl in 75% ethanol) and after overnight incubation at −20°C, samples were centrifuged at 3,500 rpm for 15 min at 4°C to remove cell debris. Insulin content was determined using the Rat High Range Insulin ELISA (Mercodia).

### Electron microscopy analysis

Freshly isolated islets were washed with PBS, fixed with 4% formaldehyde/0.1% glutaraldehyde in 0.1 M sodium cacodylate buffer (pH 7.4), and pelleted at 13,000 rpm for 5 min. The fixative was aspirated, and the cell pellet was resuspended in warm 10% gelatin (sodium cacodylate buffer). The islets were then pelleted (4000 rpm for 1 min) and the gelatin-enrobed pellet was set on ice for 30 min, post-fixed in 1% osmium tetroxide for 2 h, dehydrated in increasing concentrations of ethanol, stained with 2% uranyl acetate and embedded in agar low viscosity resin.

After trimming the resin block containing the cell pellets, 70 nm sections were cut using a Reichardt Ultracut E ultramicrotome. The sections were collected as ribbons of 3-4 sections on a 75-mesh grid (Ted Pella). Every grid was then poststained with 3% uranyl acetate in water for 10 minutes and Reynold’s lead citrate for 2 minutes. EM images were taken at 2500x magnification by a JEOL TEM1400 transmission electron microscope equipped with an Olympus SIS Quemesa 11 Mpxl camera at 80 kV. Random single sections from β cells (15 cells total) that possessed multiple dense-core granules and well-fixed cellular constituents (mitochondria, nuclear material and plasma membrane) were selected for analysis. The plasma membranes, granules and nuclear borders of each β cell were manually marked in transmission electron microscopy (TEM) micrographs. Scanned images were elaborated with the open-source software Microscopy Image Browser (MIB) to generate a mask of the cell compartments and the nucleus, and imported into ImageJ (National Institutes of Health, Bethesda, MD, USA) for image analysis.

### Cell culture and transfection

The mouse insulinoma cell line βTC3 was cultured in DMEM:F12 (1:1) supplemented with 10% heat inactivated fetal bovine serum, 100 U/ml penicillin, and 100 μg/ml streptomycin. For overexpression experiments, βTC3 cells were transfected with plasmids encoding mouse furin and/or mouse insulin receptor (pcDNA3.1 backbone) and/or mouse *Atp6ap1*/*Ac45*-Flag using Lipofectamine 2000 (Life Technologies) according to the manufacturer’s protocol. For knockdown experiments, βTC3 cells were transfected with the SMARTpool siGENOME mouse Insr siRNA (Dharmacon, cat. #16337), *Atp6ap1/Ac45* (Dharmacon cat. #M-060210-01-0005) or *Atp6ap2/Prr* (Dharmacon cat. #M-063641-02-0005) using Lipofectamine RNAiMax (Life Technologies) according to the manufacturer’s protocol. The Stealth siRNA Negative Control, Med GC (Life Technologies) or the siGENOME non-targeting siRNA control pools (Dharmacon cat. #D-001206-13-05) was transfected as a control. Cells were harvested 48 hours after transfection. Insulin receptor and PRR knockdown were verified by Western blotting, Atp6ap1/*Ac45* knockdown was confirmed by qPCR. For insulin stimulation experiments, βTC3 cells (~80% confluency) were starved overnight in serum-free DMEM:F12 medium containing 0.1% bovine serum albumin (BSA), and penicillin-streptomycin. Subsequently, cells were cultured for 2 h in glucose-free modified KRB buffer (125 mM NaCl, 4.74 mM KCl, 1 mM CaCl_2_, 1.2 mM KH_2_PO_4_, 1.2 mM, MgSO_4_, 5 mM NaHCO_3_, 25 mM HEPES [pH 7.4]) with 0.1% BSA to minimize endogenous insulin secretion. The stimulation was performed in the same buffer with 100 nM insulin final concentration for 5 minutes.

### Generation of Furin knockout βTC3 cells using the CRISPR-Cas9 nuclease system

A furin knockout βTC3 cell line was generated using the CRISPR-Cas9 nuclease system, following the protocol as described before (61). Briefly, three 20-nucleotide guide sequences targeting *fur* exon 2 were designed using the online CRISPR Design Tool (http://crispr.mit.edu). The pSpCas9(BB)-2A-Puro construct (Addgene 48139) was used to clone the guide sequences. Optimal transfection and puromycin selection conditions for βTC3 cells were established in advance using a GFP-containing expression construct. To screen guide efficacy, transfected cells were harvested for DNA extraction using the QuickExtract solution, and indel mutations were detected by the SURVEYOR nuclease assay as described (61). Clonal cells were obtained by serial dilution in 96-well plates (Corning). Genomic microdeletions and insertions were verified by TOPO cloning and subsequent Sanger sequencing. Furin deficiency in βTC3 cells was corroborated by western blot using an anti-furin antibody. Sequences for guide cloning in Addgene vector 48139, and SURVEYOR primer sequences are shown in **Table S9**.

### Apoptosis assay

Apoptosis was measured in freshly isolated islets using the Cell Death ELISA kit (Roche) according to the manufacturer’s protocol.

### Cell proliferation assay

βTC3 cells were plated in triplicates at a density of 1 × 10^4^ cells/well in 96-well plates and then incubated for 72 h. Cell proliferation was measured using the mitochondrial dye WST-1. Briefly, cells were incubated with 10 μl of WST-1 reagent per well for 2h, and the absorbance was measured at 450 and 600 nm. Three independent experiments were performed with the results presented ± SEM for the mean of all assays.

### Western blot

Freshly isolated islets were lysed in 2X Lysis buffer (Cell Signaling Technology) supplemented with protease and phosphatase inhibitor cocktails (Roche) by sonication on ice. βTC3 or HEK293T cells were lysed in 1X Ripa and extraction lysis buffer (Thermo Scientific) supplemented as above. Protein concentration was determined by BCA analysis (Thermo Scientific). Samples were boiled for 10 minutes in 4% β-mercaptoethanol and 1x sample buffer (Thermo Scientific) and loaded on a 10% Bis/Tris gel with MES or MOPS running buffer for SDS-PAGE analysis. Proteins were transferred to a nitrocellulose blot, blocked with blocking buffer (5% non-fat milk, 0.2% Triton X-100 in PBS) and subsequently incubated with primary antibody in blocking buffer at 4°C overnight. Blots were washed in PBS 0.2% Triton X-100, incubated with peroxidase-conjugated secondary antibody (Dako) for 1h, and proteins were detected using the Western Lightning ECL System (PerkinElmer). For deglycosylation experiments, 100 μg of proteins from the cell lysate were boiled at 95°C for 5 min in sodium phosphate buffer 0.1% SDS and 2-Mercaptoethanol (diluted 1:6 in water) in a total volume of 35ul. Once cooled down, the boiled samples were supplemented with 0.8% NP-40 and 1 Unit of N-glycosydase F (Roche) in a final volume 50ul and incubated at 37°C overnight. The samples were then processed for western blot as above. The primary antibodies used were rabbit anti-ATF4, anti-phospho-p70S6K (Thr389), anti-p70S6K, anti-phospho-PERK (Thr980), anti-PERK, anti-phospho-4E-BP1 (Thr37/46), anti-4E-BP1, anti-phospho-IRS1 (Ser318), anti-IRS1, anti-Akt, anti-phospho-Akt (Ser473), and anti-GAPDH from Cell Signaling Technology; mouse anti-CHOP and, anti-eIF2α from Cell Signaling Technology; rabbit anti-phospho-IRE1α (Ser724) and anti-Renin R from Novus Biologicals; mouse anti-flag M2 from Sigma; mouse anti-ATF6α from Novus Biologicals; mouse anti-furin and rabbit anti-phospho-eIF2α (Ser51) were generated as previously described (62, 63). Mouse anti-insulin receptor was a gift from Dr. Ken Siddle (University of Cambridge).

To determine secreted sPRR levels in culture medium, proteins were precipitated with methanol. Briefly, βTC3 cells were plated in 6-well plates at 80% confluency, washed and incubated overnight at 37°C and 5% CO_2_ in 1ml of serum-free DMEM:F12 medium. Sixteen hours after incubation the conditioned medium was transferred to ice cold tubes and centrifuged at 100xg at 4°C for 10 minutes. Then, an equal amount (12.5μg) of bovine serum albumin (BSA) was added to each supernatant. After the addition of BSA, the samples were vigorously vortexed and 4 ml of cold methanol was added and shaken by hand. After a 2 hour incubation at −20°C, the samples were centrifuged at 1500 xg at 4°C for 15 minutes. Then, the supernatants were removed and the pellets were dried for 2 hours. Dried pellets were dissolved in 25 μl of sample buffer 2X and analyzed by Western blot as described above.

### Microarray analysis

Microarray analysis was performed as described before (22). Briefly, islets were isolated from male Flox and βfurKO mice (n=4 per group, 12 weeks of age) by collagenase injection in the pancreatic duct. Islet RNA was isolated using an Absolutely RNA Microprep Kit (Stratagene) according to the manufacturer’s protocol. RNA quantity and quality were determined using a bioanalyser (2100; Agilent). Microarray analysis was performed on MoGene_1.0_ST arrays (Affymetrix). One hundred ng of total islet RNA was hybridized to the arrays according to the manufacturer’s manual 701880Rev4. Samples were analyzed pairwise, and p<0.01 and fold change ≥ 1.5 were set as selection criteria. Functional enrichment and canonical pathway enrichment were performed using Ingenuity Pathway Analysis (IPA, Qiagen). To identify upstream regulators, the IPA Upstream Regulator Analysis tool was used.

### Quantitative RT-PCR (qRT-PCR)

RNA from mouse islets or βTC3 cells was isolated using the Nucleospin RNA II (Macherey Nagel) kit according to the manufacturer’s protocol. cDNA was synthetized using the iScript cDNA synthesis kit (Bio-Rad). Primers were designed using the ProbeFinder software (Roche Applied Sciences). qRT-PCR was performed in triplicate with a MyiQ single-color real-time PCR detection system (Bio-Rad) using SYBR Green. RNA from human islets (∼150 for each donor) was extracted with RNeasy Mini Kit (Qiagen) and was reverse transcribed using the High Capacity Reverse Transcription Kit (Applied Biosystems). qRT-PCR was performed with a ViiA 7 detection system from Life Technologies (Thermo Fisher Scientific) using SYBR Green PCR master mix (Bioline). Data is represented as 2^-ΔCt compared to average of PC1/3 in healthy group. Primers for human and mouse genes are listed in **Table S9**.

### Stable Isotope Labeling by Amino acids in Cell culture (SILAC)

Spike-in SILAC analysis was performed to compare relative abundance of proteins in islets from male Flox and βfurKO mice, following established protocols (64). To prepare the spike-in standard, MIN6 cells were cultured for 12 passages in SILAC medium (Dulbecco’s modified Eagle medium (DMEM), high glucose (4,5 g/l) without L-glutamine, arginine and lysine (Silantes Basic Products cat. #280001300), supplemented with 15% fetal bovine serum (FBS), glutamine PS, heavy arginine (+10) and heavy lysine (+8) (Silantes Basic Products cat. no. 282986444). Light and heavy labeled cells were lysed in freshly prepared SDT-lysis buffer (2% SDS, 50 mM DTT, 100 mM Tris HCl pH 7.6), DNA was sheared by sonication, the sample was incubated at 95°C for 5’ and centrifuged. To verify the labeling efficiency, 25 μg of the cell lysate of the light labeled cells (L) was mixed with equal amounts of heavy labeled cells (H) and digested with trypsin using the FASP protocol (64). After analysis of the sample on a Q Exactive mass spectrometer (Thermo Scientific), more than 99% of the peptides had a log_2_ normalized ratio between −1 and 1, demonstrating efficient labeling.

### Sample preparation for proteomic analysis

Islets from 8 WT and 8 βfurKO mice were isolated and were pooled per 2 mice of the same genotype to obtain at least 20 μg of proteins upon lysis in SDT-lysis buffer. Twenty μg of islet lysate was pooled with equal amounts of the spike-in standard and digested with trypsin using the FASP protocol (64). Using strong anion exchange (SAX) on a StageTip format (Thermo Scientific), 3 fractions (pH 3, 6 and 11) were obtained and analyzed on a Q Exactive mass spectrometer. The quality of the spike-in standard was analyzed as described before (64). For more than 90% of the proteins the difference in experimental sample and the standard was <5, suggesting that the spike-in standard is of good quality.

### Mass spectrometry

Samples were analyzed by LC-MS/MS, coupling an Easy nLC1000 nanoflow HPLC system to the Q Exactive benchtop mass spectrometer (both Thermo Fisher Scientific) as described previously (65). Data were analyzed using the MaxQuant software environment (version 1.3.0.5) as previously described. Statistical evaluation of the data was performed with the freely available Perseus statistics software (version 1.2.0.7) and Microsoft Excel. Common contaminants and reverse decoy matches were removed from the protein identification list. At least 2 unique peptides per protein were required for protein identification. H/L ratios were reversed to L/H ratios. Only proteins that were identified and quantifiable in at least three biological replicates in each group were used for relative quantification. The average group means of the ratios were calculated and data are expresses as the log2 value of the ratio of ratios (βfurKO/control). For statistical evaluation, a two-sided t-test was used. The p-value was corrected using false discovery rate (FDR) based multiple hypothesis testing. Both t-test and FDR based multiple hypothesis testing were carried out with the default settings of the Perseus statistics software.

### Statistical analysis

Results are expressed as means ± SEM. Statistical analysis was performed by unpaired Student’s t test or two-way ANOVA with post-hoc Bonferroni correction for pairwise time-specific differences between genotypes. GraphPad Prism 8 software was used for all analyses. A value of p<0.05 was considered significant. *p<0.05, **p<0.01, ***p<0.001.

## Author Contributions

BB, BRM and JWC designed the research. BB, IC, and BRM performed the majority of *in vivo* and cell culture experiments. BB, KV and IC performed the EM analysis. NJ and JLE performed the experiments with human islets. BD, SFL and JD performed the proteomics analysis. LVL and FS performed the microarray analysis. BB, LT and JD performed bioinformatics analysis of microarray and proteomics data. BB, IC, BRM and JWC collected and analyzed the data and wrote the manuscript with input from the rest of co-authors. All the authors approved the final version of the manuscript.

## Acknowledgements

We would like to thank Sandra Meulemans, Elisa Cauwelier, Charlotte Segers and Cindy Baldwin for technical assistance, and to Dr. Jens Brüning (Max Planck Institute for Metabolism Research) for providing the IR-floxed mice. This work was supported by the FWO Vlaanderen (grant number G0B0617N) and by the Deutsche Forschungsgemeinschaft (DFG, German Research Foundation) under Germany’s Excellence Strategy within the framework of the Munich Cluster for Systems Neurology (EXC 2145 SyNergy-ID 390857198). JLE is a Chercheur-boursier from the FRQS and her research was supported by a grant from the CIHR (PJT-148771). BRM was supported by the Miguel Servet Type I program (CP19/00098) from the Institute of Health Carlos III (ISCIII), and co-financed by the European Regional Development Fund. BB and IC were supported by predoctoral fellowships from the FWO Vlaanderen.

## Supplementary Figures

**Figure S1.**
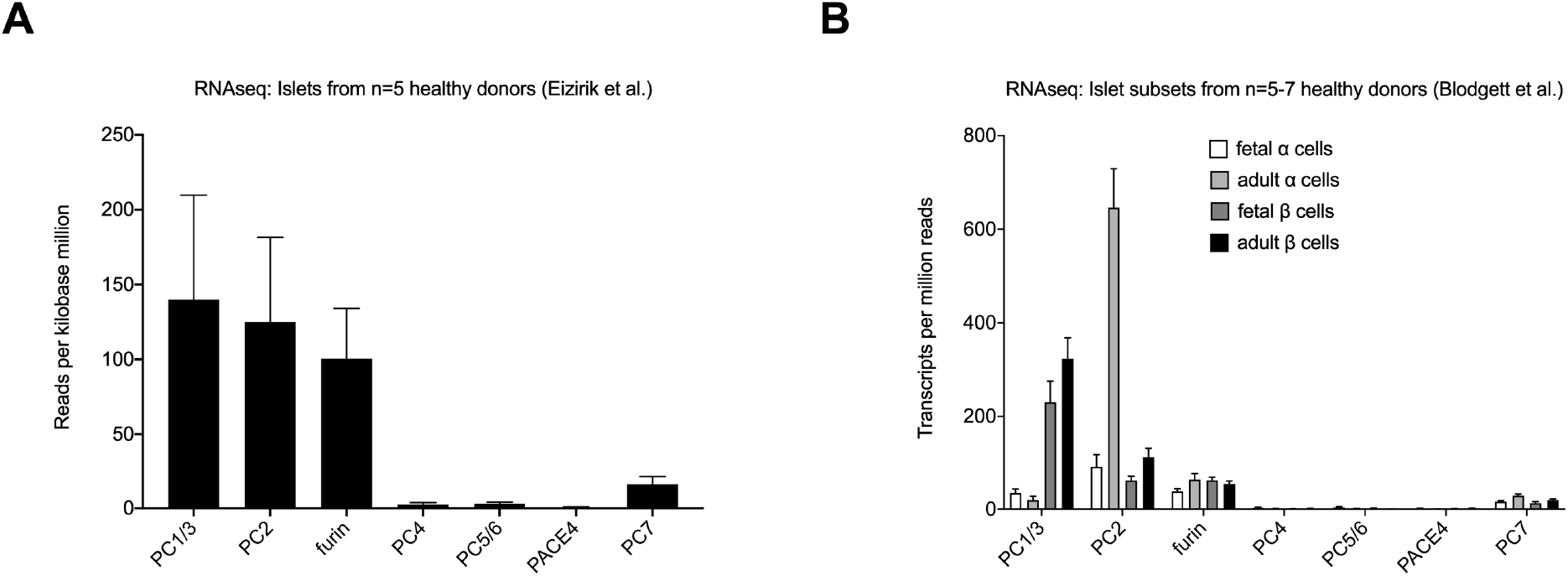
(A) RNA sequencing data of islets from healthy human donors (n=5) (taken from (19)). Data are shown as reads per kilobase million. (B) RNA sequencing data from islet subsets (i.e. fetal or adult α or β cells) from 5-7 healthy donors (taken from (20)). **Related to Figure 1**.

**Figure S2.**
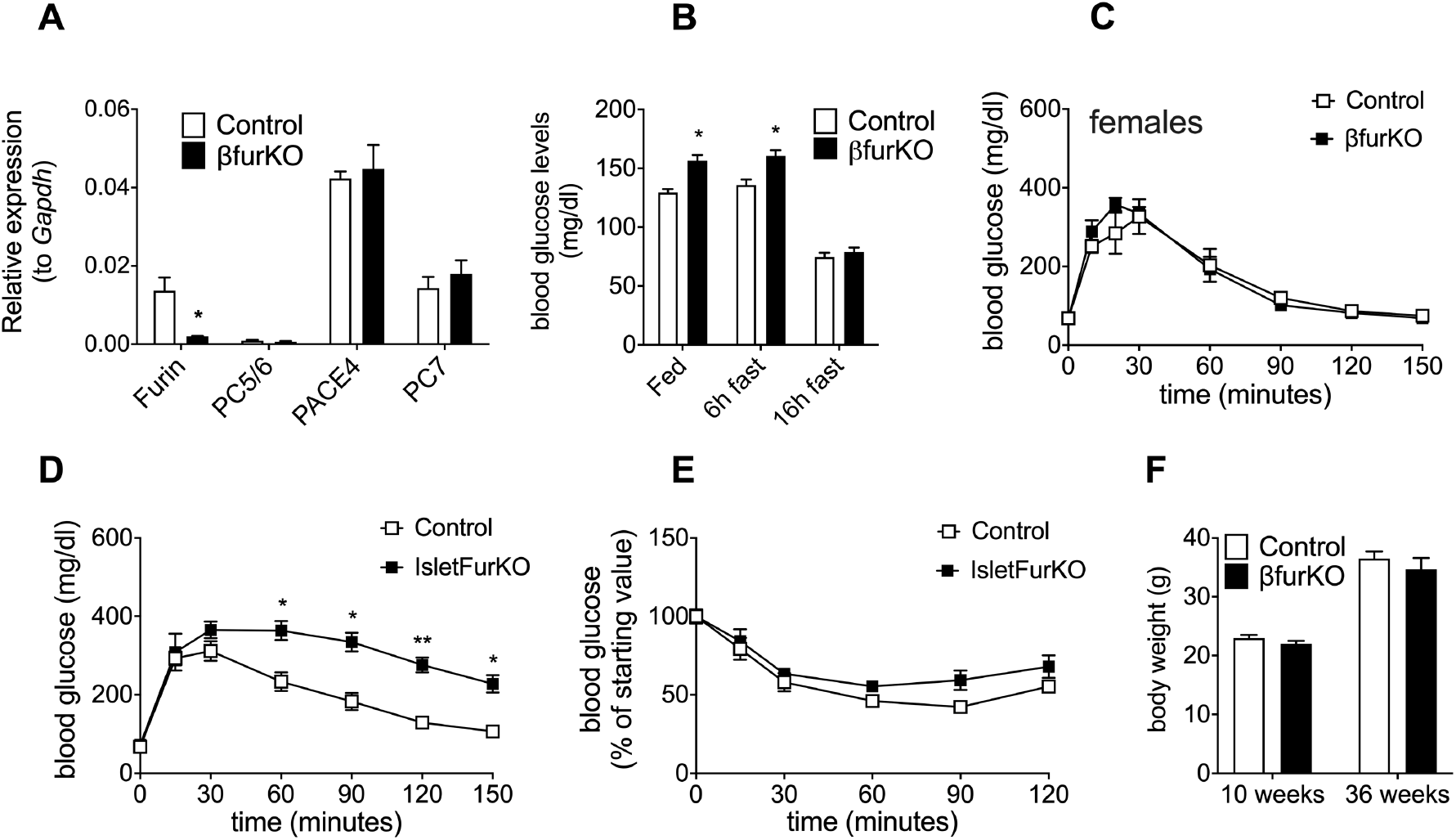
(A) qRT-PCR analysis of islet *Furin*, *PC5/6, PACE4*, and *PC7* mRNA from 12-week-old βfurKO and control mice, n=3-5 per group, **P<0.01. (B) Blood glucose levels measured in 10-week-old βfurKO and control mice, in a random fed state, and after a 6h or an overnight (ON, ~16h) fast, n=12-23 animals per group, *P<0.05. (C) IPGTT on 10-week-old female mice, (n=4)/genotype. (D) IPGTT and (E) IPITT on male Ngn3-Cre+/−; fur^flox/flox^ (‘IsletFurKO’) and control mice at 36 weeks of age, n=4-6 per group. (F) Body weight (g) of 10-week-old and 36-week-old control and βfurKO mice (n=9-10/group). **Related to Figure 2**.

**Figure S3.**
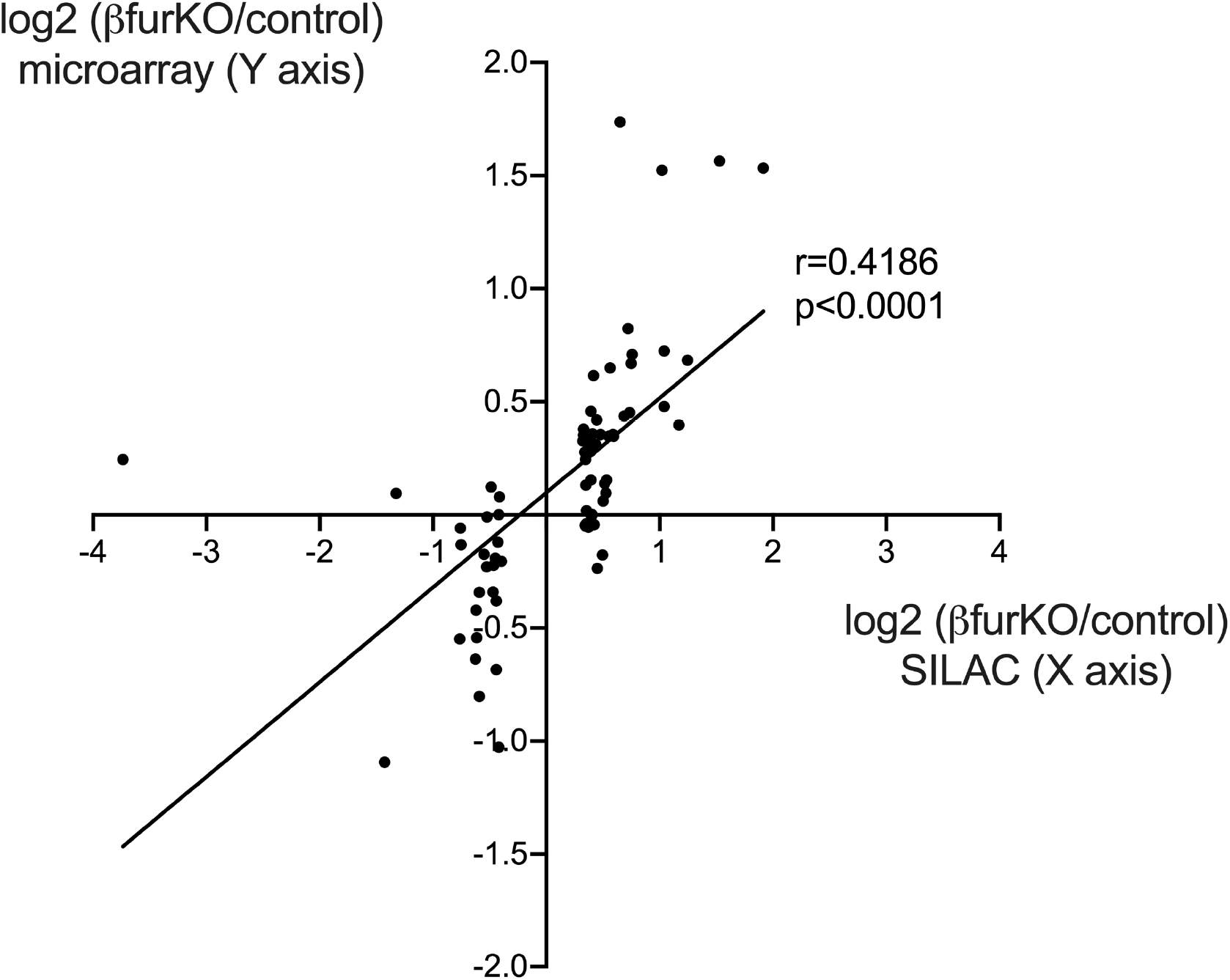
Correlation between SILAC and microarray data (r=0.4186, p<0.0001), for proteins that show at least a 25% increase or decrease in expression as quantified in the SILAC experiment. **Related to Figure 4**.

**Figure S4.**
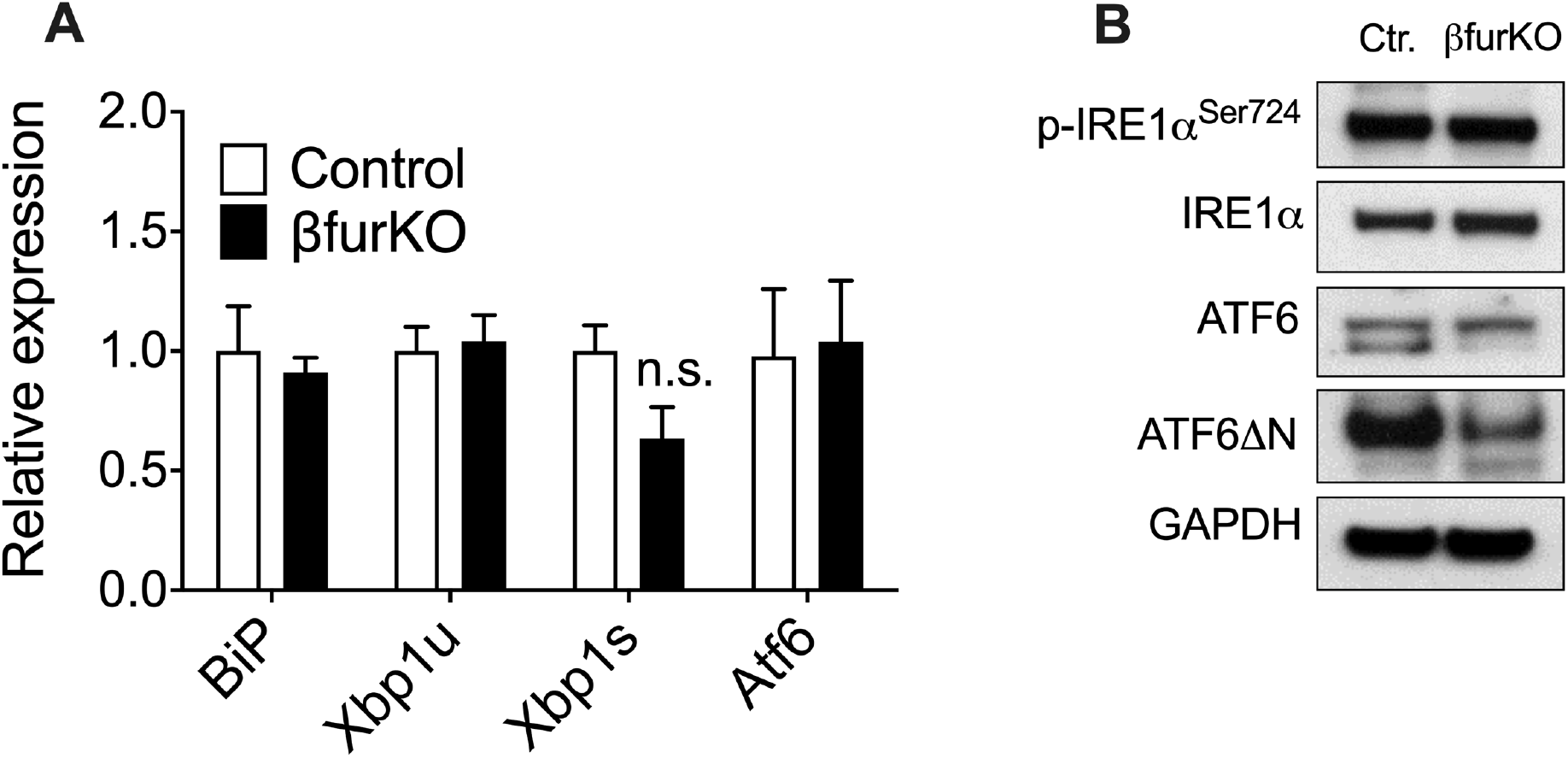
(A) qRT-PCR expression of genes involved in the unfolded protein response (UPR): binding immunoglobulin protein (*BiP*), unspliced and spliced X-box binding protein 1 (*Xbp1u*, *Xbp1s*) and Activating transcription factor 6 (*Atf6*), n=3-6 per group. (B) Protein analysis of total and phosphorylated Inositol-requiring protein 1 (IRE1α), and uncleaved and cleaved activating transcription factor 6 (ATF6, ATF6ΔN). GAPDH was used as a loading control. **Related to Figure 5**.

**Figure S5.**
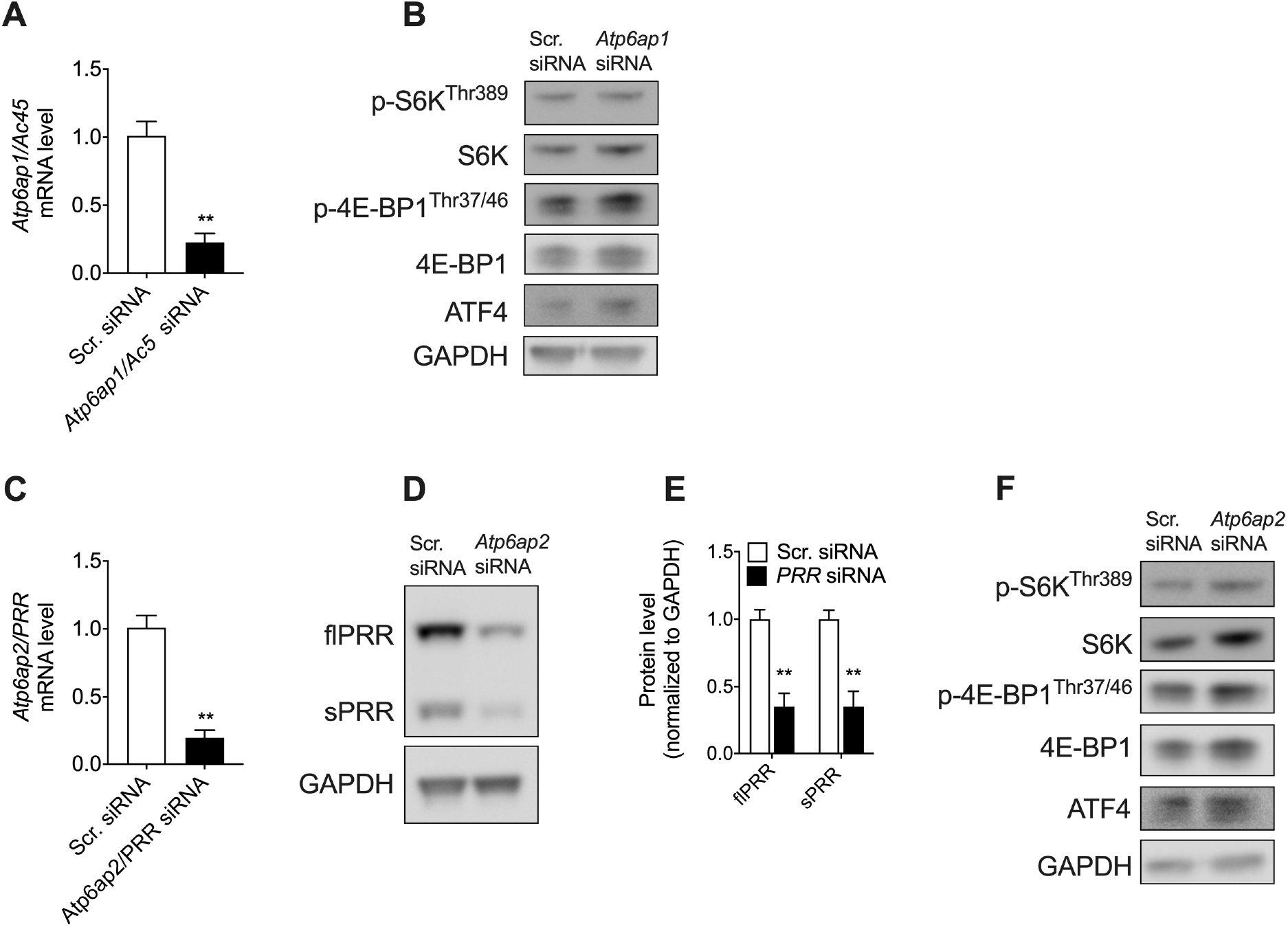
(A) Knockdown efficiency of *Atp6ap1/Ac45* in βTC3 cells measured by qRT-PCR, n=3 independent experiments. (B) Representative western blot of phosphorylated and total levels of S6K and 4E-BP1, and total levels of ATF4 in βTC3 cells upon knockdown of *Atp6ap1/Ac45*. GAPDH was used as a loading control. (C-D) Knockdown efficiency of *Atp6ap2/Prr* in βTC3 cells measured by qRT-PCR (C) and western blot (D), including western blot quantification (E), n=3 independent experiments. (F) Representative western blot of phosphorylated and total levels of S6K and 4E-BP1, and total levels of ATF4 in βTC3 cells upon knockdown of *Atp6ap2/Prr*. GAPDH was used as a loading control. **P<0.01. **Related to Figure 7**.

